# Modeling With Uncertainty Quantification Identifies Essential Features of a Non-Canonical Algal Carbon-Concentrating Mechanism

**DOI:** 10.1101/2024.04.12.589284

**Authors:** Anne K. Steensma, Joshua A.M. Kaste, Junoh Heo, Douglas J. Orr, Chih-Li Sung, Yair Shachar-Hill, Berkley J. Walker

## Abstract

The thermoacidophilic red alga *Cyanidioschyzon merolae* survives its challenging environment likely in part by operating a carbon-concentrating mechanism (CCM). Here, we demonstrated that *C. merolae*’s cellular affinity for CO_2_ is stronger than its rubisco affinity for CO_2_. This provided further evidence that *C. merolae* operates a CCM while lacking structures and functions characteristic of CCMs in other organisms. To test how such a CCM could function, we created a mathematical compartmental model of a simple CCM distinct from those previously described in detail. The results supported the feasibility of this proposed minimal and non-canonical CCM in *C. merolae*. To facilitate robust modeling of this process, we incorporated new physiological and enzymatic data into the model, and we additionally trained a surrogate machine-learning model to emulate the mechanistic model and characterized the effects of model parameters on key outputs. This parameter exploration enabled us to identify model features that influenced whether the model met experimentally-derived criteria for functional carbon-concentration and efficient energy usage. Such parameters included cytosolic pH, bicarbonate pumping cost and kinetics, cell radius, carboxylation velocity, number of thylakoid membranes, and CO_2_ membrane permeability. Our exploration thus suggested that a novel CCM could exist in *C. merolae* and illuminated essential features necessary for CCMs to function.

**Significance:** Carbon-concentrating mechanisms (CCMs) are processes which boost photosynthetic efficiency. By developing modeling approaches to robustly describe CCMs in organisms where biochemical data is limited, such as extremophile algae, we can better understand how organisms survive environmental challenges. We demonstrate an interdisciplinary modeling approach which efficiently sampled from large parameter spaces and identified features (e.g., compartment permeability, pH, enzyme characteristics) which determine the function and energy cost of a simple CCM. This approach is new to compartmental photosynthetic modeling, and could facilitate effective use of models to inform experiments and rational engineering. For example, engineering CCMs into crops may improve agricultural productivity, and could benefit from models defining the structural and biochemical features necessary for CCM function.

## Introduction

*Cyanidioschyzon merolae* is a red microalga found in moist environments surrounding geothermal sulfur springs. This species is extremophilic, with optimal laboratory growth conditions including low pH (∼ 2) and high temperatures (∼ 42 °C) (1, 2). *C. merolae* and other thermo-acidophilic red algae draw interest for their unique biology and simple characteristics, which position them as useful model organisms and as candidates for biotechnology applications (3–6). For example, *C. merolae* is of interest because it is one of few organisms which relies on photosynthesis in geothermal spring environments, where hot and acidic conditions restrict the availability of inorganic carbon and challenge biological carbon fixation (1, 7). Notably, organisms of acid waters can only access approximately 10 micromolar inorganic carbon, as the inorganic carbon pool at acid pH is primarily the volatile species CO_2._ In comparison, organisms of near-neutral and alkaline waters may have access to several millimolar of inorganic carbon, due to accumulation of the involatile bicarbonate (8).

*C. merolae* is thought to survive in its challenging environment in part by operating a carbon-concentrating mechanism (CCM) (9–11). CCMs boost carbon-fixation efficiency by concentrating CO_2_ around rubisco, providing ample substrate for carbon-fixation and inhibiting a competing oxygen-fixation reaction of rubisco. Evidence supporting a CCM in *C. merolae* includes measured accumulation of carbon in the cell, δ^13^C consistent with a CCM, similar growth rates under ambient and elevated CO_2_, transcriptional response of potential CCM genes to CO_2_ fluctuations, and substantial CO_2_ assimilation at low environmental CO_2_ concentrations (9–12). However, many of these indications of the CCM are not definitive: in particular, it is not known how much of *C. merolae*’s ability to assimilate CO_2_ efficiently could be explained by the affinity of *C. merolae* rubisco for CO_2_. Thus, we here provide further evidence for the CCM in *C. merolae* by demonstrating that the affinity of *C. merolae* cells for CO_2_ is better than could be explained by the affinity of *C. merolae* rubisco for CO_2_.

*C. merolae*’s CCM may be described as a “novel” or “non-canonical” CCM, as the *C. merolae* CCM must operate differently from the few CCMs which are well-characterized. Unlike algae and cyanobacteria with well-characterized CCMs, *C. merolae* is not able to take up external bicarbonate, and *C. merolae* lacks anatomy associated with the pyrenoid CCM organelle (10, 11, 13, 14). The absence of these CCM features in *C. merolae* challenges our understanding of how algal CCMs work, and presents the opportunity to define essential CCM components. We thus used mathematical modeling, informed by new experimental measurements, to explore how the *C. merolae* CCM may function.

Research on CCMs has long employed mathematical models to understand the components of functional CCMs in model cyanobacteria and algae. A particular area of interest in CCM modeling is the possibility of boosting crop productivity by engineering CCMs into crops which lack CCMs (15–18). We sought to add to the inspiration for these engineering efforts by modeling a heat-tolerant CCM with minimal components which offers unique possibilities for plant engineering (19). To draw robust conclusions about cellular characteristics which can support a CCM, we used state-of-the-art statistical methods to define the effects of model parameters on the predicted photosynthetic phenotype while limiting unwarranted *a priori* assumptions.

Some sets of model input parameters produced model outputs which met empirically-based criteria for functional carbon concentration and efficient energy usage, and we identified input parameters which have substantial impacts on the model outputs. Overall, our model of a hypothetical biophysical CCM which requires minimal enzymes and anatomical features (**Figure 1**) appears to represent a feasible CCM structure in *C. merolae*, which invites further research into the sources of environmental resilience in extremophile algae.

**Figure 1.**
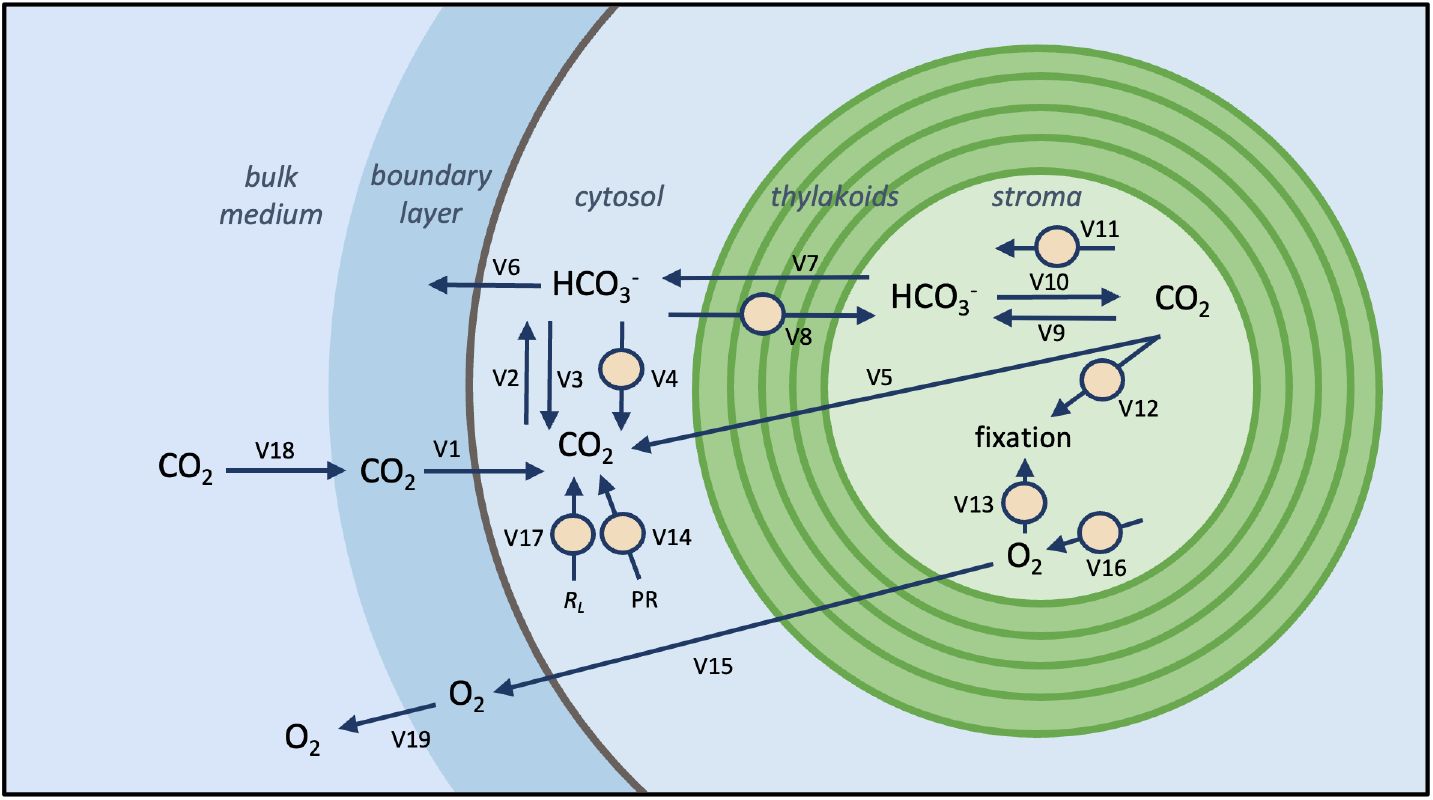
Cross-section of model structure. This model describes fluxes (indicated by arrows) and pools (indicated by molecular formulas) of a simplified dissolved inorganic carbon system (CO_2_, HCO_3_^-^) and of oxygen (O_2_). Molecule pools can be present in several well-mixed compartments: the bulk external medium surrounding the cell, an unstirred boundary layer of medium around the cell, the cytosol, or a central stromal space of the chloroplast. Circles mark enzymatically-catalyzed fluxes. Compartments are not drawn to scale. *PR* = photorespiratory CO_2_ release, *R*_*L*_ = respiration in the light. All fluxes are reversible and are assigned an arbitrary direction, except those fluxes which represent producing or consuming material.

## Methods

### Experimental data collection

Extraction, purification, and kinetic assays of *C. merolae* rubisco; and measurement of gas-exchange parameters by open-path infra-red gas analysis, were as detailed in **Supplemental Methods**.

### Model details

The hypothetical CCM described in this study (**Figure 1**) was modeled as a set of well-mixed compartments and represented as a system of ordinary differential equations (ODEs). In this minimal biophysical CCM, carbon diffuses into the cell as CO_2_, is trapped in the cytosol as bicarbonate by action of carbonic anhydrase, and is pumped into the chloroplast, where a second carbonic anhydrase provides CO_2_ around rubisco. No pyrenoid diffusion barrier is present, though we accounted for potential effects of the concentric thylakoids which are present in *C. merolae* and many other aquatic photosynthetic organisms (22).

The model geometry is based on the cellular structure of *C. merolae* as apparent in published micrographs of this alga (22–31). The modeled cell and its boundary layer form a series of concentric spherical well-mixed compartments. The cell is enclosed by a lipid bilayer of radius *Radius*_*cell*_. The cell is surrounded by a medium boundary layer of radius 2 * *Radius*_*cell*_, beyond which lies an infinite external medium. The cell contains a cytosol of radius *Radius*_*cell*_ and a chloroplast stroma space of radius 0.25 * *Radius*_*cell*_.

Molecules cross the boundary of the stroma space according to diffusion or transport equations. For flux calculations, the boundary consists of 1 to 7 lipid bilayers of negligible thickness that are evenly spaced from 0.5 * *Radius*_*cell*_ to 0.25 * *Radius*_*cell*_. This boundary structure represents the fact that the *C. merolae* chloroplast is surrounded by a chloroplast envelope and by approximately 4 to 6 thylakoids which appear as concentric circles or spirals in thin-section microscopy (22). A range of possible transport scenarios (how many membranes molecules must cross when crossing between the cytosol and stroma, and how much energy this crossing costs) are captured by varying parameters *Membranes* and *Pump*_*cost*_.

Diffusion through lipid membranes was described using estimates of conductivity of lipid membranes to the chemical species in question:

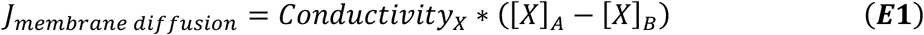

Where *Conductivity*_x_ is the conductivity – in units of μm^3^/s – of chemical species X through a lipid bilayer, and *[X]*_A_ and *[X]*_B_ are the concentrations of that species on the two sides of that lipid bilayer. Diffusion into or out of the medium boundary layer was described as an analogous simple diffusion flux, with conductivity determined according to diffusion coefficients through water at the boundary layer thickness. Lipid permeability coefficients for CO_2_ and HCO_3_^-^ and the water diffusion coefficient for O_2_ were sourced from the literature (**Table S1**), and other necessary gas permeability and diffusion coefficients were determined from the literature values by Graham’s law of diffusion:

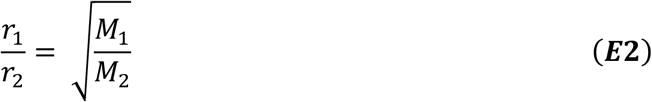

Where the rates of diffusion *r*_*1*_ and *r*_*2*_ for two different ideal gases, here CO_2_ and O_2_, are related according to their two molar masses *M*_*1*_ and *M*_*2*_.

To describe diffusion of CO_2_, HCO_3_^-^, and O_2_ through variable numbers of stacked thylakoid membranes, an overall conductivity through all of the layers was calculated as:

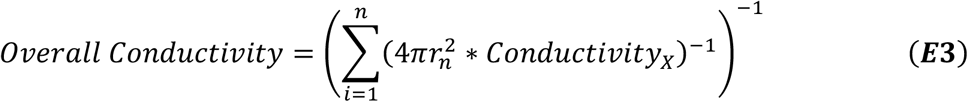

Where *r*_*n*_ is the radius of the sphere formed by the *n*th thylakoid membrane. This overall conductivity value is then used in **(E1)** to describe the movement of a chemical species from the outer stroma into the inner stroma space, as shown in **Figure 1**. We assume that small gas molecules diffuse easily around membrane proteins, so that the diffusion of CO_2_ and O_2_ through any modeled membrane is potentially impeded by increased path length, but is not impeded by CO_2_ and O_2_ passing through high-resistance protein material.

Spontaneous interconversion of CO_2_ and HCO_3_^-^ **(E4-5)** was described using simple first-order kinetics, according to the rate constant of the dehydration (slower) step of the interconversion:

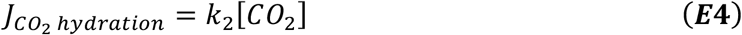

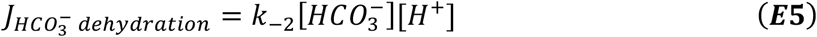

Note that CO_2_ must first be hydrated to H_2_CO_3_, which is then deprotonated to yield the HCO_3_^-^ ion. However, because the interconversion of HCO_3_^-^ and H_2_CO_3_ is essentially instantaneous relative to the hydration-dehydration reaction, here we ignore the H_2_CO_3_ species and approximate the spontaneous interconversion as the hydration-dehydration reaction.

The interconversion of CO_2_ and HCO_3_^-^ by carbonic anhydrase was described as in (16):

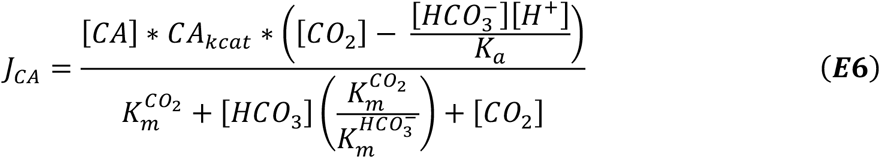

Where the K_a_ value is the overall K_a_ for the CO_2_/HCO_3_^-^ system. This value is temperature-sensitive and was calculated using the R package *seacarb* package (32). Other potentially temperature-sensitive parameters receive temperature adjustments according to Q_10_ or Q_15_ factors.

Carboxylation by rubisco was described as with the assumption that CO_2_ is limiting, as in (33):

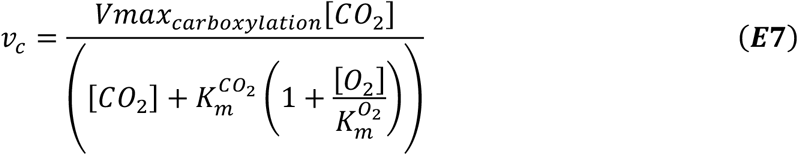

To estimate oxygenation, we estimate *v*_*c*_*/v*_*o*_ (carboxylation flux over oxygenation flux) from the CO_2_/O_2_ specificity (*S*_*c/o*_) of rubisco and chloroplast CO_2_ and O_2_ concentrations **(E8)**, and then use this to arrive at *v*_*o*_.

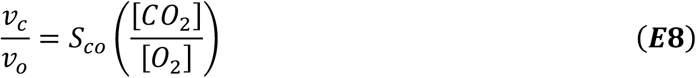

The pumping of HCO_3_^-^ across the stack of thylakoid membranes by a bicarbonate pump was described by simple Michaelis-Menten kinetics:

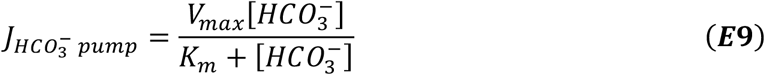

Respiration in the light (*R*_*L*_) was estimated from experimental data according to a modified Kok method, by measuring under sub-saturating light intensities and extrapolating CO_2_ release in the absence of light (**Figure 2**). The mean measured value of *R*_*L*_ was normalized to cell size for use in the model: we assume that the empirical measurement of *R*_*L*_ we obtained was, on a per cell basis, characteristic of a *C. merolae* cell of a radius of 1 μm. Under the assumption that *R*_*L*_ should vary proportionally with cell volume, we normalized *R*_*L*_ as follows:

**Figure 2.**
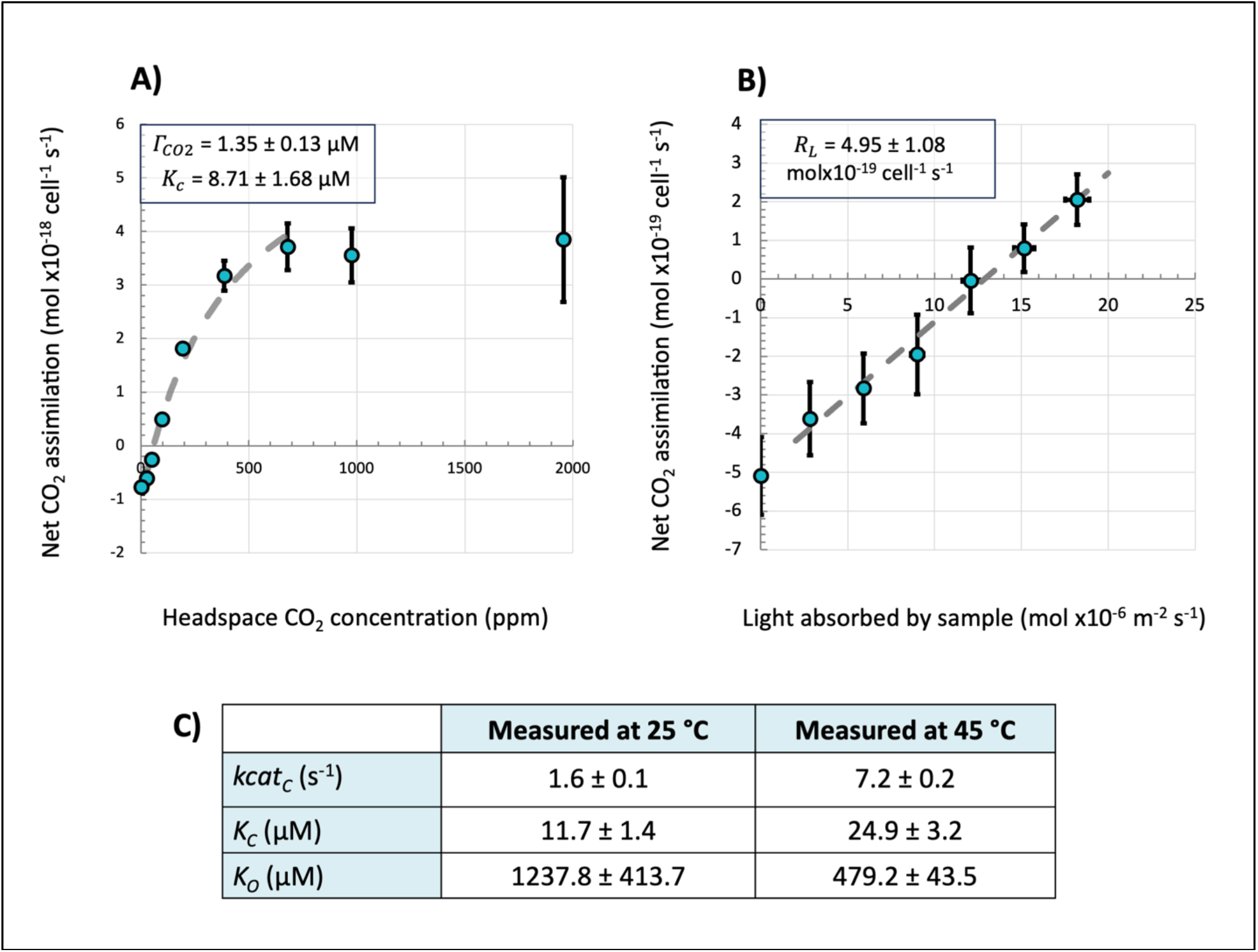
Experimental data incorporated into the model. **(A**,**B)**. Response of net assimilation in *C. merolae* to **(A)** CO_2_ availability and **(B)** light availability. Points are mean ± SE (n = 3), and parameters calculated from the data are indicated in the upper left corner of each plot as mean ± SE. Dashed lines indicate trend fits used to determine *Kc* and *R*_*L*_. The linear fit used to determine *Γ*_*CO2*_ is not pictured. **(C)** Kinetic properties of *C. merolae* rubisco. Rubisco turnover rate for CO_2_ fixation (*kcat*_*C*_), Michaelis-Menten constant of CO_2_ fixation (*K*_*C*_), and Michaelis-Menten constant of O_2_ fixation (*K*_*O*_) were measured at 25 and 45 °C. Data is mean ± SE, n = 4.

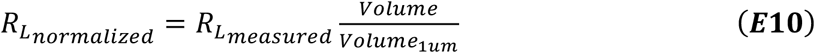

ATP costs for the cell were estimated as:

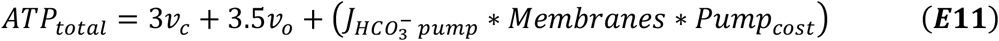

Where *Membranes* is the number of thylakoid stacks and *Pump*_*cost*_ is the assumed cost, in ATP, of pumping a single HCO_3_^-^ ion across a lipid bilayer by the hypothesized pump.

A full list of all flux equations and the system of ODEs used to describe the system can be found in **Supplemental Materials**.

### Definition of reasonable model output values

To ensure the model reproduced experimental results, we used new and published experimental data to set acceptable bounds for the following model outputs: CO_2_ compensation point (*Γ*_*CO2*_), the ratio of ATP consumption flux to net CO_2_ assimilation flux (ATP per CO_2_), the steady-state CO_2_ concentration in the chloroplast stroma (stromal CO_2_), and the ratio of oxygen-fixation flux to carbon-fixation flux (*v*_*o*_*/v*_*c*_). Selection and justification of these bounds are detailed in **Supplemental Methods**.

### Model optimization and estimation of simulated compensation point

Steady-state fluxes and metabolite concentrations were solved using *odeint()* from Python’s SciPy library (34). Latin hypercube parameter sampling (35) and curve-fitting to generate compensation point estimates were as detailed in **Supplemental Methods**.

### Parameter exploration and surrogate model selection

In order to thoroughly explore the 19-dimensional parameter space in a computationally-feasible way, we trained a surrogate machine-learning model on the mechanistic CCM model. By emulating the intricacies of the mechanistic model, surrogate modeling faithfully captures dynamics of complex systems while alleviating the substantial computational costs associated with obtaining results. Surrogate modeling additionally gave us access to powerful statistical tools for machine-learning model analysis, including SHapley Additive exPlanations (SHAP) (36) and partial dependence (PD) plots (37).

To identify the optimal surrogate model for parameter exploration, we compared four popular machine-learning models: eXtreme Gradient Boosting (XGBoost) (38), Local approximate Gaussian Process (laGP) (39), single-layer Neural Network (NN) (40), and Deep Neural Network (DNN) (38). We collected a 240,000-sized dataset, where the outputs were simulated from the CCM model at space-filling input locations. 90% of the data was used for training the surrogate, and the remaining 10% was used as the test dataset to validate the model performance. The evaluation of prediction performance was based on the root-mean-square error (RMSE):

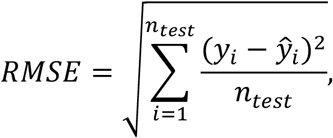

where *y*_*i*_ is the *i*-th test output and *ŷ*_*i*_ is the *i*-th predicted model output.

Model outputs had varying scales and degrees of skew, so to effectively compare prediction performance on different model outputs, a normalized RMSE (NRMSE) was calculated. The NRMSE was calculated as the RMSE divided by 4_*max*_ − 4_*min*_, where 4_*max*_ is the highest test output and 4_*min*_ is the lowest test output.

From the model evaluation (**Table S2**), it appears that XGBoost outperformed other models for *v*_*o*_*/v*_*c*_ and ATP per CO_2_, and remained comparable for *Γ*_*CO2*_ and stromal CO_2._ As such, XGBoost was used as the surrogate model for further analyses.

## Results and Discussion

### Rubisco kinetics demonstrated that *C. merolae* operates a CCM

In previous work, we determine that if *C. merolae* has rubisco kinetics similar to other red algae, then this alga must operate a CCM to maintain its measured photosynthetic efficiency. Alternatively, its measured photosynthetic efficiency could be explained by unprecedented rubisco kinetics, meaning enzyme properties favoring carbon-fixation over oxygen-fixation to an unprecedented degree (10). Here we confirmed that *C. merolae* rubisco kinetics are similar to those of other red-type (Form 1D) rubiscos (41–43). *C. merolae* rubisco had a strong affinity for CO_2_ (low *K*_*C*_), a poor affinity for O_2_ (high *K*_*O*_), and a slow carboxylation rate (low *kcat*_*C*_) (**Figure 2**). Consistent with other studies, *kcat*_*C*_ and *K*_*C*_ were higher when measured at increased temperature, while *K*_*O*_ was lower. Although *K*_*O*_ is a component of rubisco specificity (*S*_*c/o*_) and *S*_*c/o*_ decreases with increased temperature, *in vitro K*_*O*_ is observed to decrease with increased assay temperature in some species (42, 44, 45).

These kinetics findings indicated *C. merolae* does operate a CCM, as *C. merolae* cells had higher affinity for CO_2_ than *C. merolae* rubisco (8.71 ± 1.7 μM cell *K*_*C*_ vs. 24.9 ± 3.2 μM rubisco *K*_*C*_ at 45 °C, *p* = 0.008 by two-sample *t*-test) (**Figure 2**). This result adds to the indications of the CCM in *C. merolae* (9–11).

### Quantitative modeling showed that a hypothesized CCM can explain *C. merolae*’s carbon-concentrating behavior

To explore how the *C. merolae* CCM may operate, we constructed a functional model of a CCM (**Figure 1**). This model demonstrated that there were parameter sets consistent with the empirical literature that result in a functional CCM, despite the minimal model structure (**Figure 3**). Our results provided quantitative support for a CCM taking inorganic carbon from the environment solely through CO_2_diffusion into the cell, which we term a “non-canonical” or “novel” CCM due to its differences in structure and function from CCMs that have been characterized in detail. Though there is speculation that extremophilic red algae may use a C_4_-like CCM, it has been previously proposed that acidophile algae may accumulate carbon by a “bicarbonate-trap” or “acid-loading” mechanism similar to our modeled CCM (7, 12, 15, 46, 47). Briefly, bicarbonate would be concentrated for enzymatic action by bringing inorganic carbon speciation near equilibrium in near-neutral cellular compartments, since the predominant inorganic carbon species from pH ∼6 to ∼10 is the poorly-membrane-permeable bicarbonate.

**Figure 3.**
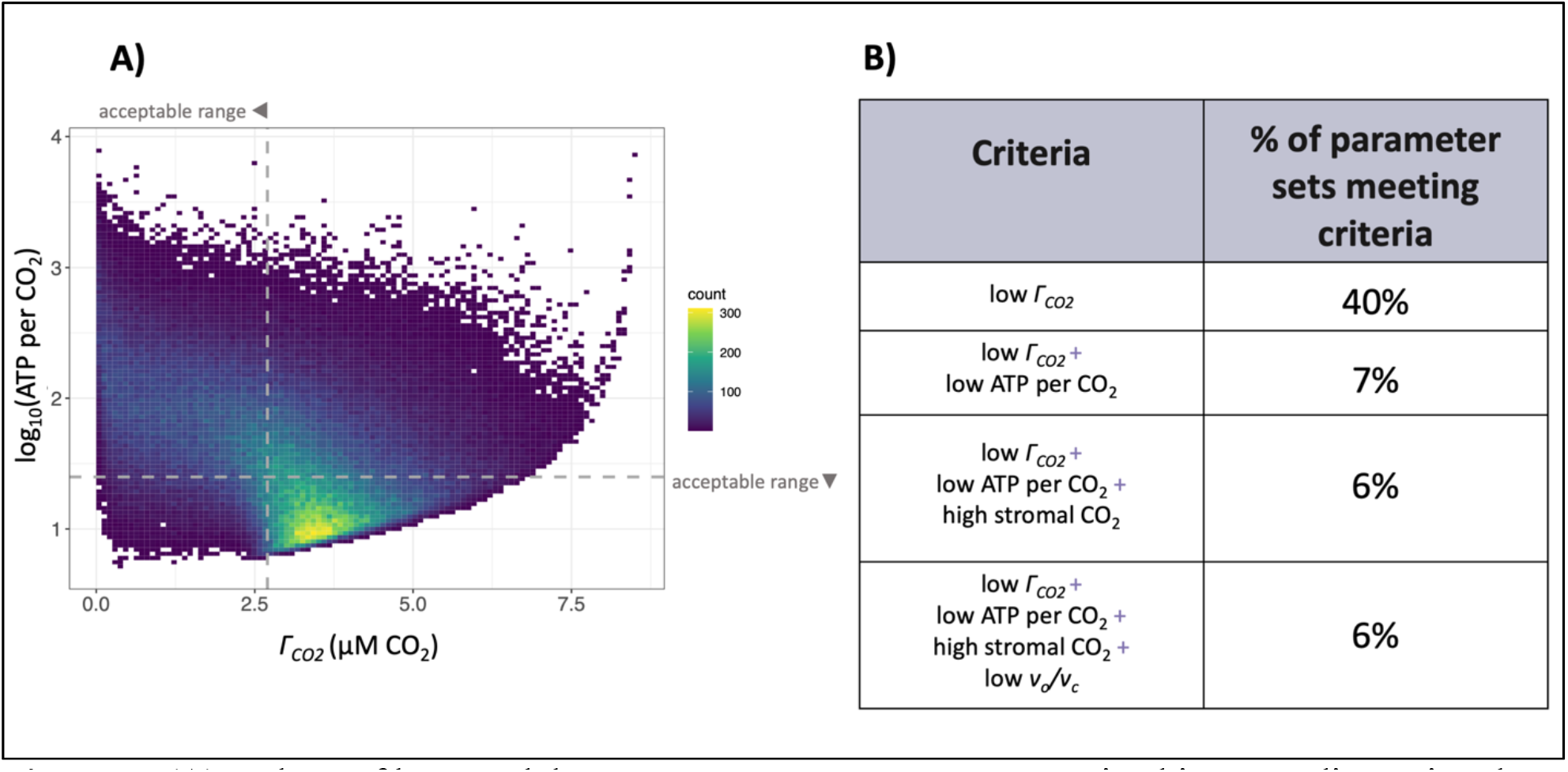
**(A)** Values of key model outputs. Parameter sets are organized into a 2-dimensional histogram according to their output values of *Γ*_*CO2*_ and ATP per CO_2_, with dashed lines indicating bounds for acceptable values of these outputs. 80 parameter sets (0.03% of total) are not pictured on the figure, as they produced negative ATP per CO_2_ values and could not be log-transformed. **(B)** Percentages of parameter sets meeting various combinations of output criteria.

We used two strategies to deeply explore the model parameter space and ensure that our conclusions were robust. First, the model included new experimental data on gas-exchange and rubisco parameters central to photosynthetic efficiency (**Figure 2**). Second, we developed a method for thoroughly assessing the model’s sensitivity to the value of model parameters of interest. Specifically, we were interested in 19 of the 43 model parameters which were biologically interesting in relation to the function of a novel CCM and which were not well-characterized physical constants (**Table S1**). We thus sampled input parameter sets through a Latin hypercube design (35). Latin hypercube sampling enhanced analysis accuracy by mitigating sampling bias, as it produced parameter sets distributed throughout the 19-dimensional parameter space of interest. Then, each input parameter set was used to parameterize the model and to generate a set of outputs for analysis.

Some of the input parameter sets produced outputs consistent with a functional CCM with reasonable energy cost. Of particular interest were the parameter sets which met all the empirically-based criteria for a realistic and functional CCM (criteria selection described in **Supplemental Methods**). 13,998 of 240,000 (6%) of parameter sets fulfilled the two competing objectives of functional carbon concentration (corresponding to outputs of low *Γ*_*CO2*_, high stromal CO_2_, and low *v*_*o*_*/v*_*c*_) and efficient energy usage (corresponding to output of low ATP per CO_2_) (**Figure 2**).

The generated parameter sets allowed us to explore the trade-offs associated with various features related to the CCM. For example, adding additional concentric thylakoids slightly improved carbon concentration by presenting barriers to CO_2_ leakage out of the chloroplast, but incurred additional energy costs (**Figure 4, Figures S1 – S2**). This is consistent with other modeling studies indicating that thylakoid membranes could affect inorganic carbon diffusion (15, 48).

**Figure 4.**
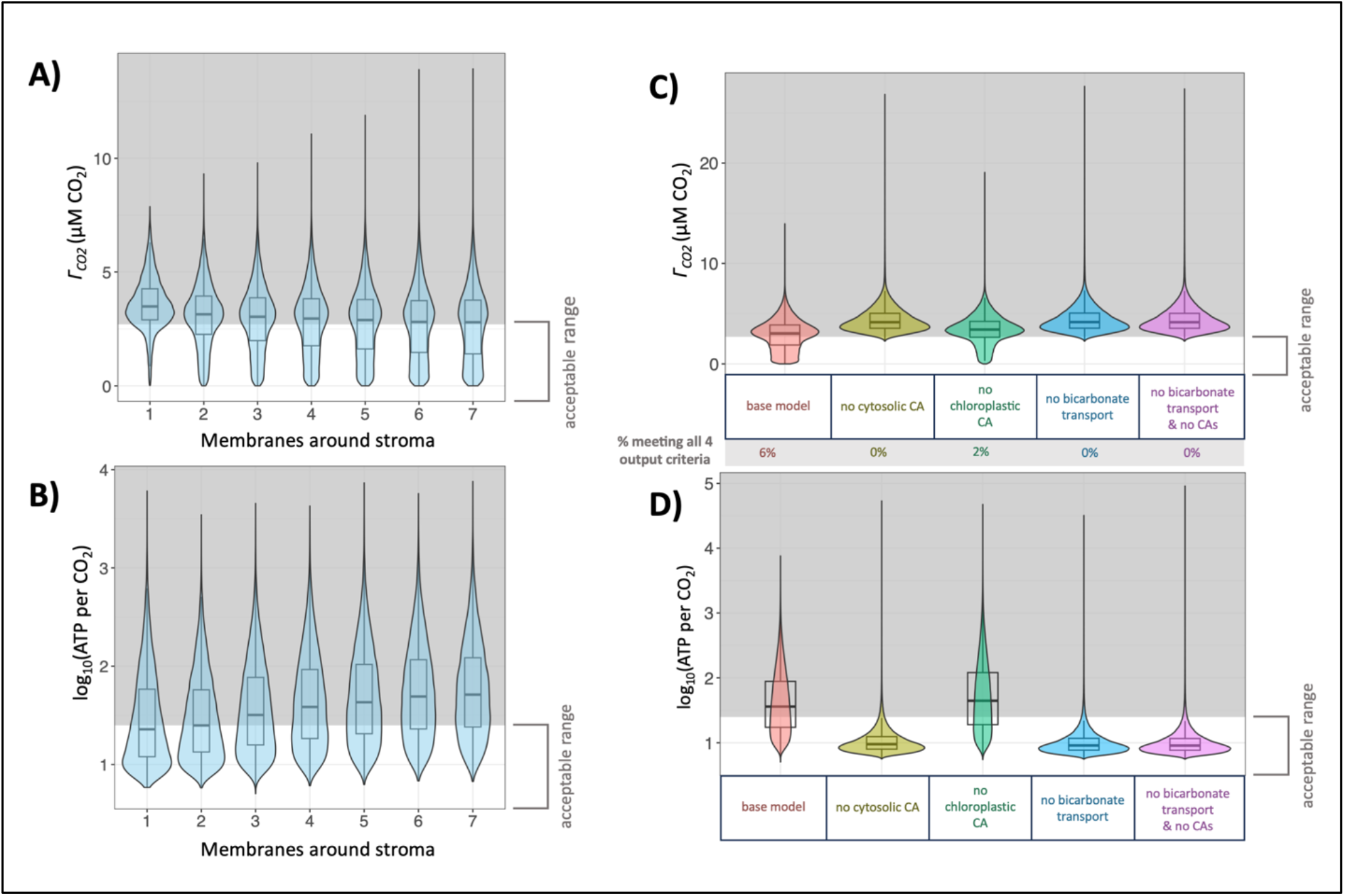
**(A, B)** Effect of model input parameter *Membranes* (x-axis) on key model outputs. Distribution of parameter set outputs for each value of *Membranes* is represented by a box plot overlaid on a violin plot. Shaded areas represent unacceptable values of outputs. **(A)** Effect of *Membranes* on model output *Γ*_*CO2*_. **(B)** Effect of *Membranes* on model output ATP per CO_2_. 80 parameter sets (0.03% of total) are not pictured in this panel, as they produced negative ATP per CO_2_ values and could not be log-transformed. **(C, D)** Effect on key model outputs when bicarbonate transport or carbonic anhydrases (CAs) are removed from the model. Distribution of parameter set outputs for each scenario is represented by a box plot overlaid on a violin plot. Shaded areas represent out-of-bounds values of outputs. The same sampling of input parameter sets was run through models representing each scenario. **(C)** *Γ*_*CO2*_ in model scenarios where various model features removed, with indication of how many parameter sets met output criteria in each scenario. **(D)** ATP per CO_2_ in model scenarios where bicarbonate transport activity at the chloroplast boundary is removed. 6,991 parameter sets producing negative ATP per CO_2_ values (0.6% of total) are not pictured in this panel.

### Machine-learning-based surrogate models identified the parameters that most influence CCM efficiency

Like most mathematical models of photosynthetic systems, this model faced the challenge of drawing robust conclusions while using parameters which, although bounded by their relationship to physical processes, have substantial uncertainty (**Table S1**). To model a system with limited biochemical data while not constraining input parameters to a greater degree than was supported by the literature, it was important to assess uncertainties which seemed likely to have substantial and interdependent effects on the model. For example, the input parameter describing permeability of a lipid bilayer to CO_2_ (*Plip*_*CO2*_) has reported values ranging over several orders of magnitude (**Table S1**). Furthermore, the effect of *Plip*_*CO2*_ in the model depended on the value of other parameters, such as the number of lipid bilayers which pose a barrier to carbon moving between the stroma and cytosol (*Membranes*). *Plip*_*CO2*_ and similar parameters were unlikely to be satisfactorily explored by classical local sensitivity analyses, which involve tracking model outputs when individual parameters are varied by a set fraction of the parameter’s original value. Therefore, to reveal which model conditions were necessary for the modeled CCM and to identify interesting directions for future investigation, we used statistical methods to identify impactful parameters and to identify which input spaces corresponded to target output ranges. These statistical methods involved training a surrogate machine-learning model on our CCM model inputs and outputs. Interpretations of this surrogate model identified which zones in the input parameter space contained the most combinations fulfilling output criteria **(Figure 5 lower left)**, quantified how much each input parameter affected the prediction of outputs by the surrogate model (**Figure 5 upper right**), and visualized the response of model outputs to inputs (**Figures S4 – S7**).

**Figure 5.**
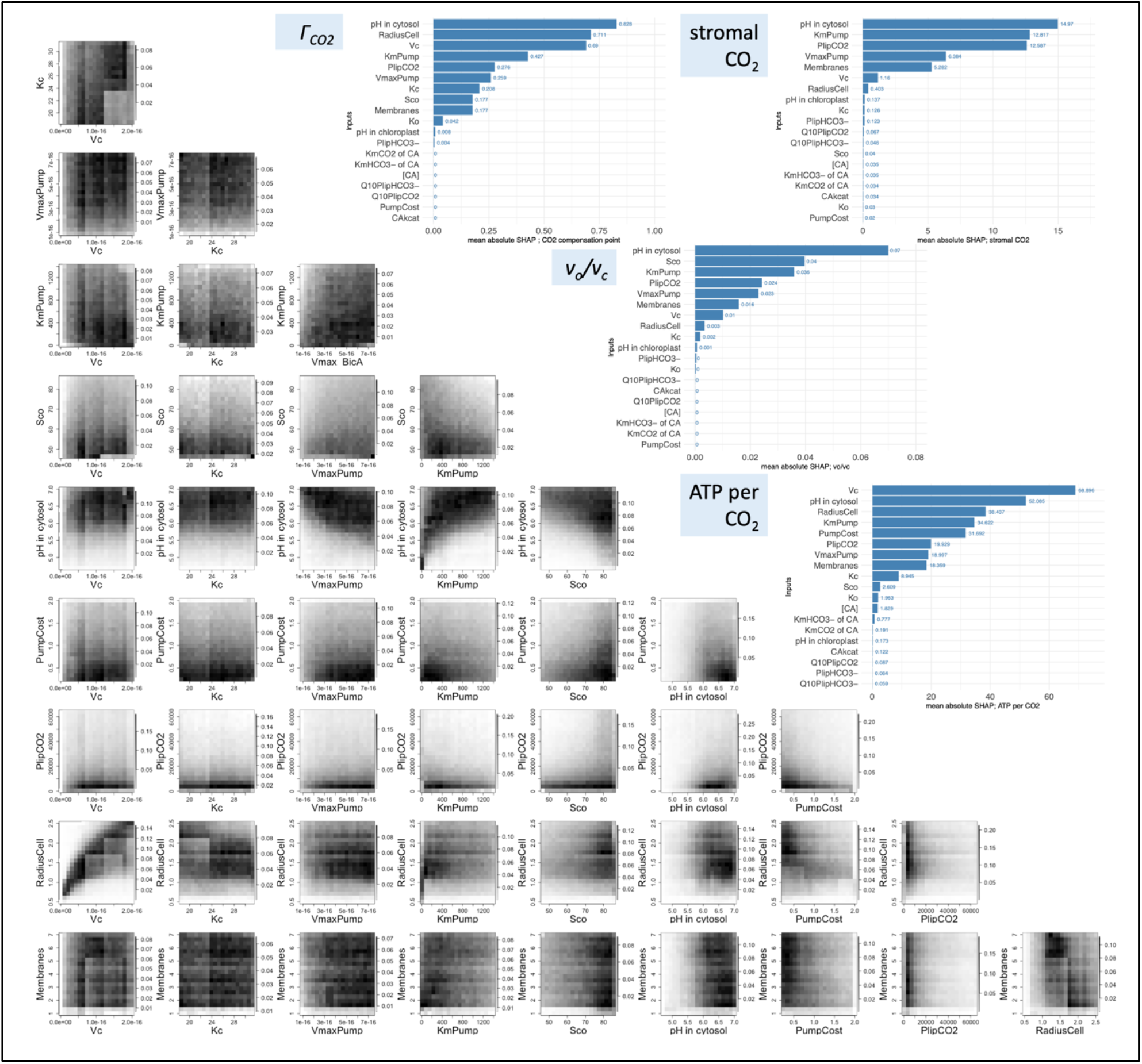
**(upper right bar plots)** Mean absolute SHapley Additive exPlanations (SHAP) plots for each output criterion. **(lower left density plots)** Density plots of parameter sets meeting all output criteria, organized by selected pairwise input parameter (input parameters pictured are those input parameters with high SHAP values for all output criteria). Darker areas indicate areas where more parameter sets meeting criteria occur. Scales of color vary for each plot).

Some input parameters had little impact on model outputs. For these parameters, values from across the input range were evenly represented in the parameter sets meeting all output criteria, which is reassuring for future modeling and engineering efforts that involve these features. The parameters with relatively little impact on outputs included values related to carbonic anhydrase concentration and kinetics (*[CA], CAkcat, Km*_*CO2*_ and *Km*_*HCO3-*_ for carbonic anhydrases), chloroplast pH, and values related to bicarbonate membrane permeability (*Plip*_*HCO3-*,_ *Q10*_*PlipHCO3-*_, **Figure 5, Figures S4 – S8**).

Other parameters were more constraining in the model, indicating their importance in producing a functional CCM. For example, six parameters appeared to impact all four of the target model outputs in the mean absolute SHAP plots: *V*_*c*_, *Vmax*_*pump*_, *Km*_*pump*_, pH in the cytosol, *PlipCO*_*2*_, and *Membranes*. As might be expected in a model relying on a cytosolic bicarbonate trap followed by bicarbonate pumping, parameter sets that successfully and efficiently concentrated carbon tended to have cytosolic pH at or above the pH where bicarbonate predominates (cytosol pH above 6), and tended to have a lower ATP cost of pumping bicarbonate (low *Pump*_*cost*_), as well as faster and higher-affinity bicarbonate pumps (high *Vmax*_*pump*_, low *Km*_*pump*_).

Other features enriched in parameter sets meeting output criteria were a cell radius in the middle of the input range (moderate *Radius*_*cell*_), and a lower CO_2_ membrane permeability (low *Plip*_*CO2*_, **Figure 5, Figures S4 – S9**). This suggested an important relationship between the volumes where metabolism occurs and the surface areas which present diffusion barriers between compartments. As the radius of the cell increases, CO_2_ loss from *R*_*L*_ may overcome the ability of the cell to acquire carbon through passive diffusion into the cell. Conversely, as the radius of the cell decreases, less absolute bicarbonate pumping would be necessary to achieve high rubisco saturation, especially when rubisco is slow (low *V*_*c*_). In low-radius scenarios, “over-pumping” bicarbonate could reduce energy efficiency.

### *In silico* knockouts identified experimental targets for further characterization of the *C. merolae* CCM

The modeling also suggested interesting directions for investigating enzymatic components of the CCM. Alternative models with CCM enzymes removed (carbonic anhydrases or bicarbonate pumping not functional) were less likely to meet the criterion of a *Γ*_*CO2*_ indicative of functional carbon concentration, but tended to have lower ATP per CO_2_ cost than the model with all enzymes present (**Figure 4, Figures S1 – S2**).

The modeled CCM functioned without fine details of cellular structure that support photosynthesis in other organisms, such as rubisco aggregation into an area smaller than the stroma, recapture of mitochondrially-respired CO_2_, and perforations or interconnections in concentric thylakoids (9, 49, 50). It may still be of interest to explore whether similar structures exist in *C. merolae*, and to investigate the biochemical and molecular basis for this novel CCM.

### Further applications of surrogate modeling and uncertainty quantification

More broadly, the statistical approach adopted in this paper represents an advance in metabolic and biochemical modeling. By training a surrogate model on the parameter space of mechanistic biological models, we can understand and account for high-dimensional uncertainty in model parameters. Metabolic modeling in general has been highlighted as a particularly promising application of surrogate modeling, since metabolic modeling has biotechnological potential but is challenged by the complexity of metabolism and by the “trial and error” process which is often required to produce a working metabolic model (21). Surrogate modeling has found uses in dynamic flux balance analysis and process modeling for bioprocesses (51, 52). Our work expands on these investigations by demonstrating what is to our knowledge the first application of surrogate modeling to ODE-based compartmental modeling of biological systems. Our methods may be particularly valuable for models that have poorly-defined parameters or are extremely computationally expensive. For example, the implementation of surrogate modeling described here could alleviate current limitations in interpreting reaction-diffusion models and genome-scale metabolic models (21).

Effective parameter exploration and analysis may generally be useful in confronting global challenges. Here, we used statistical sampling, surrogate modeling, and uncertainty quantification methods to investigate how aquatic organisms achieve the high photosynthetic efficiency that enables them to be responsible for approximately half of global photosynthetic CO_2_ consumption (53). Similar modeling techniques may be applied effectively to any system: for example, as part of engineering efforts for bioproduction, crop resilience, and other goals, it may be useful to determine which features of a system are essential or inflexible *in silico* before devoting resources to *in vivo* experimentation.

In conclusion, the extremophilic red microalga *C. merolae* operates a CCM, as evidenced by this alga having gas-exchange behavior which was not explained by its rubisco properties. Mathematical modeling suggested that this CCM could consist of a minimal mechanism which includes thylakoid membranes as diffusion barriers. Robust parameter exploration and statistical analysis, aided by the use of a surrogate model, allowed us to quantify the sensitivity of our model to parameter uncertainties, identify important parameter interactions, and identify key determinants of CCM efficiency. Therefore, in addition to supporting the presence of a novel CCM in *C. merolae*, our results shed light on what conditions must be met for this CCM to function and the essential elements of biophysical CCMs in general.

## Code availability

Model code used in this study can be accessed via GitHub: https://github.com/anne-steensma/Cmerolae_CCM_model.

## Acknowledgments and funding sources

Work in the laboratory of BJW is supported by Grant Number DE-FG02-91ER20021 from the Division of Chemical Sciences, Geosciences and Biosciences, Office of Basic Energy Sciences of the United States Department of Energy. Work in the laboratory of YSH was supported by Grant Number DE-SC0018269 from the United States Department of Energy. AKS and JAMK were additionally supported by a predoctoral training award from Grant Number T32-GM110523 from the National Institute of General Medical Sciences of the National Institutes of Health. JAMK received additional support from the National Science Foundation Research Traineeship Program, grant number DGE-1828149. JH and C-LS are supported by Grant Number DMS-2113407 from the National Science Foundation. The contents of this publication are solely the responsibility of the authors and do not necessarily represent the official views of the funding agencies.

We thank Dr. Mark Seger and Dr. Peter Lammers (Arizona Center for Algae Technology and Innovation) for kindly providing inocula and biomass of *C. merolae*. We additionally thank the following individuals for valuable supplies and technical insight: Dr. Sigal Lechno-Yossef and Damien Sheppard (Fast Protein Liquid Chromatography technical consultation), Ludmila Roze (protein gel technical consultation), Dr. Josh Vermaas and Dr. Daipayan Sarkar (insightful discussion of lipid membrane permeability).

## SUPPLEMENTAL MATERIAL

### Supplemental Methods

#### Model optimization and estimation of simulated compensation point

In order to characterize the response of key outputs and robustness of conclusions to a wide range of possible parameterizations of the model, we used Latin Hypercube Sampling to explore 240,000 parameter combinations according to the bounds specified in (**Table S1**). These simulations were run on Michigan State University’s High Performance Computing Cluster. Compensation point estimates were generated for every parameter set by running the model at external CO_2_ concentrations ranging from 0.0001 to 1000 μM, constructing a cubic spline from the resulting curve of net CO_2_ assimilation vs. external CO_2_ concentration, and identifying the root of this spline to find the compensation point. Each simulation was verified to reach steady-state (metabolite concentration solutions changing 0.01% or less from previous value).

#### Definition of reasonable output values

##### CO_2_ compensation point (Γ_CO2_)

We accepted *Γ*_*CO2*_ values less than or equal to 2.70 μM, corresponding to no more than twice the mean measured value (**Figure 2**).

##### Ratio of ATP consumption flux to net CO_2_ assimilation flux (ATP per CO_2_)

We accepted ATP per CO_2_ values which were less than or equal to 25 and greater than 0. Measured light response curves indicated how much additional light absorption drives additional CO_2_ assimilation (**Figure 2**) We used this data to estimate how much additional ATP production drives an additional CO_2_ assimilation, using the photon per ATP values for various light-reaction pathways (53), the cylindrical geometry of the gas-exchange sample chamber, and the measured density of cells in the sample. The resulting estimated values were: 13.8 ± 2.19 ATP produced/CO_2_ assimilated (mean ± SE, assuming cyclic and linear electron flow operating equally) or 17.4 ± 2.76 ATP produced/CO_2_ assimilated (mean ± SE, assuming linear electron flow only operating). This suggests that ATP per CO_2_ values of up to roughly 25 are supported by photosynthetic electron flow. The lower bound of the acceptable range excludes a few parameter sets outputting negative ATP per CO_2_, since these parameter sets represent particularly non-functional CCM scenarios with negative net assimilation values under ambient CO_2_ conditions.

##### Steady-state CO_2_ concentration in the chloroplast stroma (stromal CO_2_)

We accepted chloroplast CO_2_ concentration values of greater than or equal to the CO_2_ concentration in the medium under 400 ppm CO_2_ atmosphere, by the logic that a functional CCM should result in rubisco accessing a greater CO_2_ concentration than is available from ambient medium.

##### Ratio of oxygen fixation flux to carbon fixation flux (v_o_/v_c_)

We accepted *v*_*o*_*/v*_*c*_ values less than or equal to 0.3, based on data and models indicating that plants without CCMs are unlikely to achieve *v*_*o*_*/v*_*c*_ less than approximately 0.3 (54).

#### Experimental data collection: gas-exchange measurements

*Cyanidioschyzon merolae* 10D was grown as cultures in Erlenmeyer flasks in 50 mL of medium containing 40 mM (NH_4_)_2_SO_4_, 4 mM MgSO_4_ × 7H_2_O, 8 mM KH_2_PO_4_, 0.75 mM CaCl_2_ × 2H_2_O, 1 mL L^-1^ Hutner’s Trace Elements solution, and H_2_SO_4_ to pH 2.7 (recipe modified from MA2 medium recipe of (55)). Cultures were maintained at 40 °C under 100 μmol m^-2^ s^-1^ white light, with aeration by shaking at 100 rpm. For gas-exchange measurements, cultures of OD_750_ 1.0 – 1.2 were resuspended in growth medium to OD_750_ 0.6 (1.60×10^7^ – 3.68×10^7^ cells/mL). Gas-exchange parameters were measured in a LI-6800-18 Aquatic Chamber (LI-COR Biosciences) at 45 °C, following the procedures of (10).

#### Experimental data collection: rubisco kinetics measurements

We purified rubisco from *C. merolae* biomass with a protocol adapted from (2, 56). Approximately 60 grams of biomass were lysed by freeze-thawing followed by mechanical homogenization. Crude rubisco was polyethylene-glycol-precipitated from clarified homogenate and purified by FPLC. FPLC fractions eluting under the major UV trace peak were assayed by SDS-PAGE and by spectrophotometric rubisco activity assay (procedures adapted from (57, 58)) (**Figure S3**). Fractions containing active semi-pure rubisco were pooled, concentrated with a 100 kDa centrifugal concentration filter, and snap-frozen for use in rubisco assays.

Purified rubisco was used to determine catalytic properties as described previously (43), with some alterations to protein desalting and activation: concentrated protein aliquots were first diluted with activation mix containing 100 mM Bicine-NaOH pH 8.0, 20 mM MgCl_2_, 10 mM NaHCO_3_, and 1 % (v/v) Plant Protease Inhibitor cocktail (Sigma-Aldrich, UK). Rubisco was then activated at 45 °C for 15 min before being used in ^14^CO_2_ consumption assays at either 25 °C or 45 °C with CO_2_ concentrations of 8, 16, 24, 36, 68, and 100 μM. To determine *K*_*O*_, these CO_2_ concentrations were combined with concentrations of either 0, 21, 40, or 70 % (v/v) O_2_. *kcat*_*C*_ was determined using measurements with 0% O_2_. An aliquot of the activated protein was used for determination of Rubisco active sites via ^14^C-CABP binding using the method of (59). For ^14^C-CABP binding, protein aliquots were incubated at 45°C for 15 mins with ^14^C-CABP to maximize binding, prior to application to Sephadex columns as previously described (60). Aliquots were also analyzed via SDS-PAGE alongside known concentrations of plant type Rubisco to strengthen estimates of Rubisco content.

#### ODE System

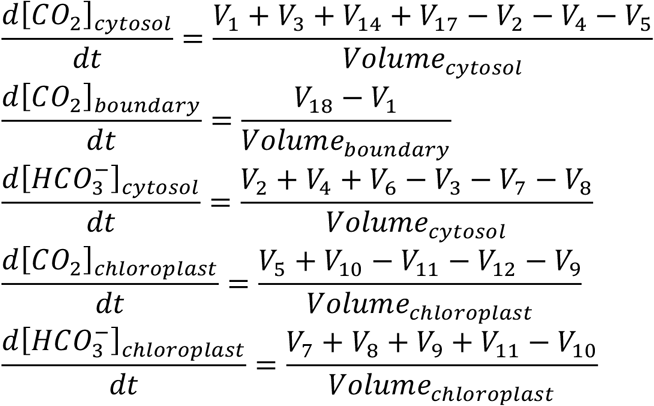

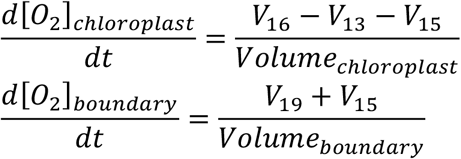

#### Model fluxes

See **Table S1** and main text for parameter sources, values, and definitions.

Diffusion of inorganic carbon through membranes or boundary layer (V1, V5, V6, V7, V15, V18, V19)

Implemented as described in Methods.

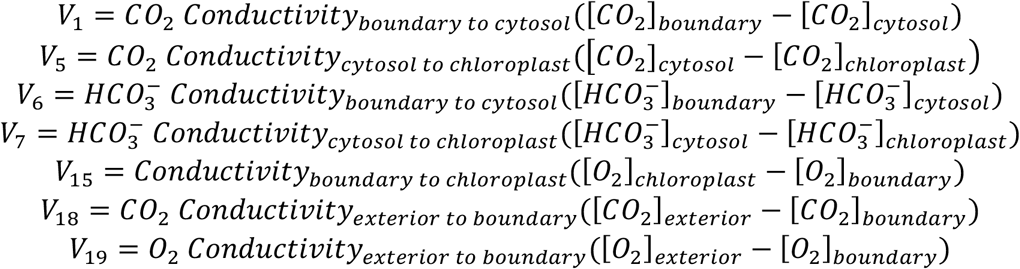

Spontaneous interconversion of dissolved inorganic carbon species (V2, V3, V9, V10)

Implemented as described in Methods.

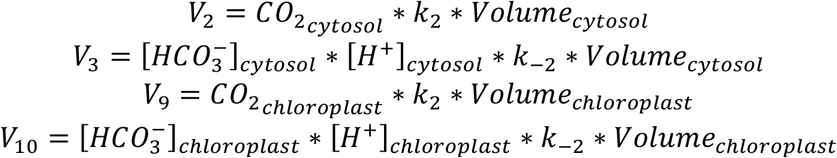

Carbonic-anhydrase-mediated interconversion of inorganic carbon (V4, V11)

Implemented as described in Methods.

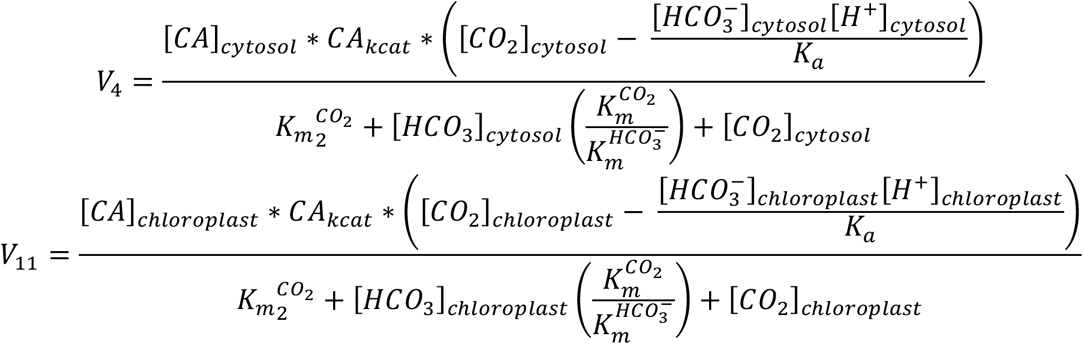

Active transport (pumping) of bicarbonate from cytosol to stroma (V8)

Implemented as described in Methods.

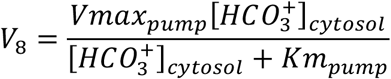

Carbon dioxide fixation by rubisco (V12)

Implemented as described in Methods.

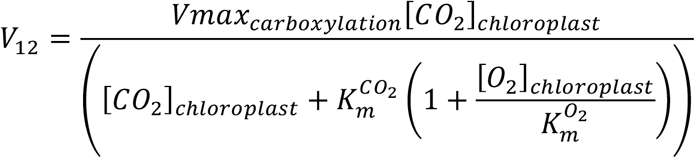

Oxygen fixation by rubisco (V13)

Implemented as described in Methods.

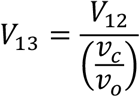

Evolution of carbon dioxide in cytosol as a result of photorespiration (V14)

This flux is determined based on the stoichiometry of photorespiration.

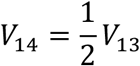

Evolution of oxygen into stroma from thylakoid action (V16)

This flux is determined based on the stoichiometry of photosynthesis.

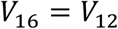

Evolution of carbon dioxide in cytosol as a result of respiration in the light (V17)

Implemented as described in Methods.

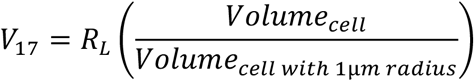

## Supplemental Figures

**Table S1.**
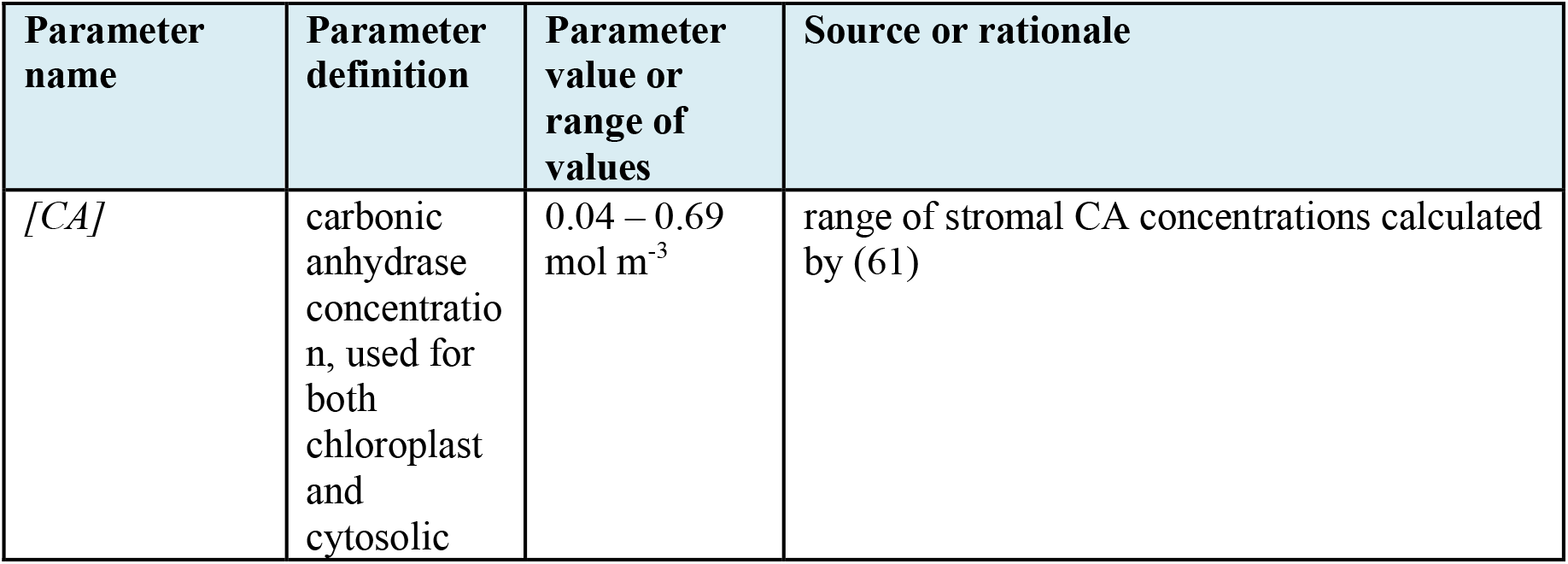
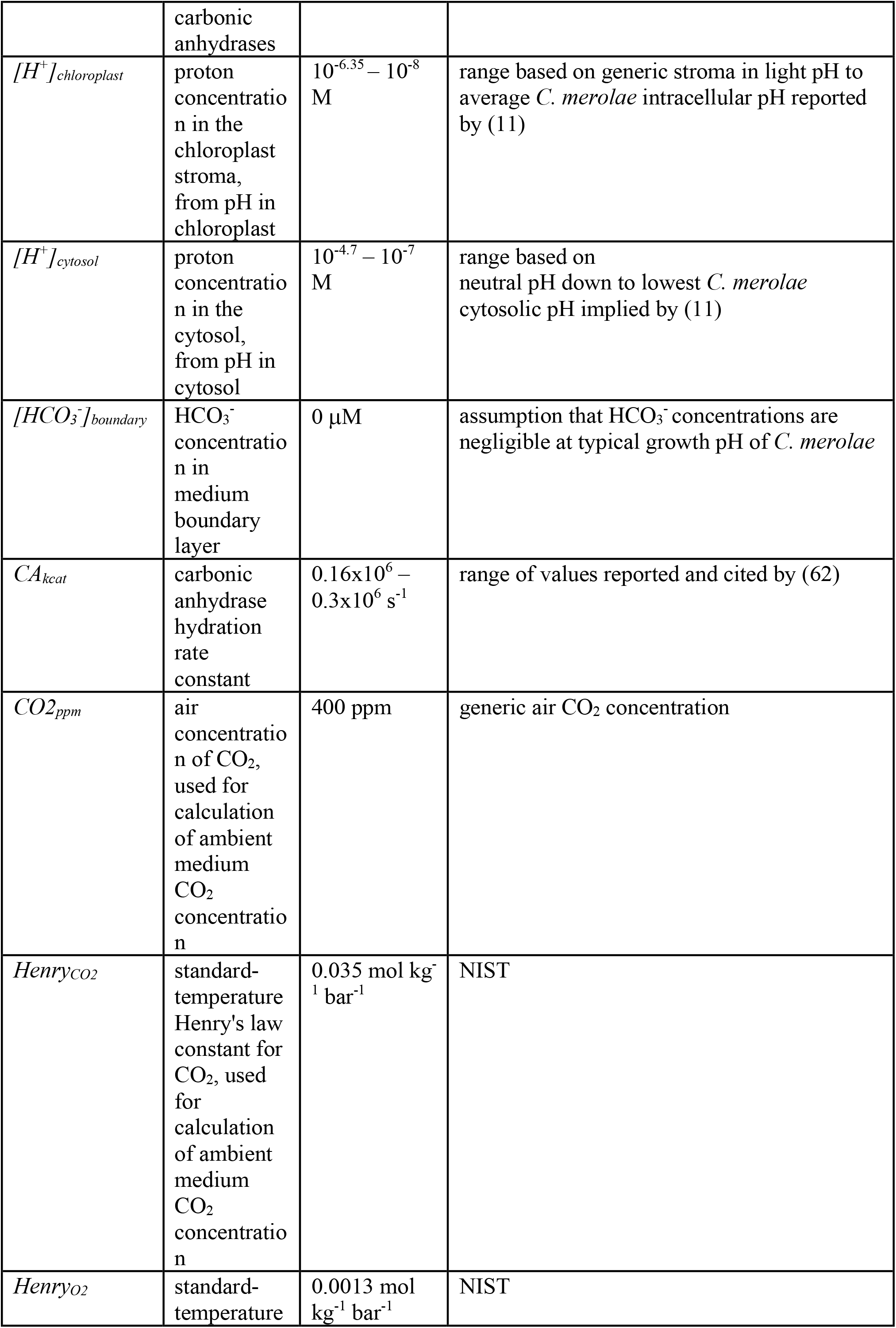
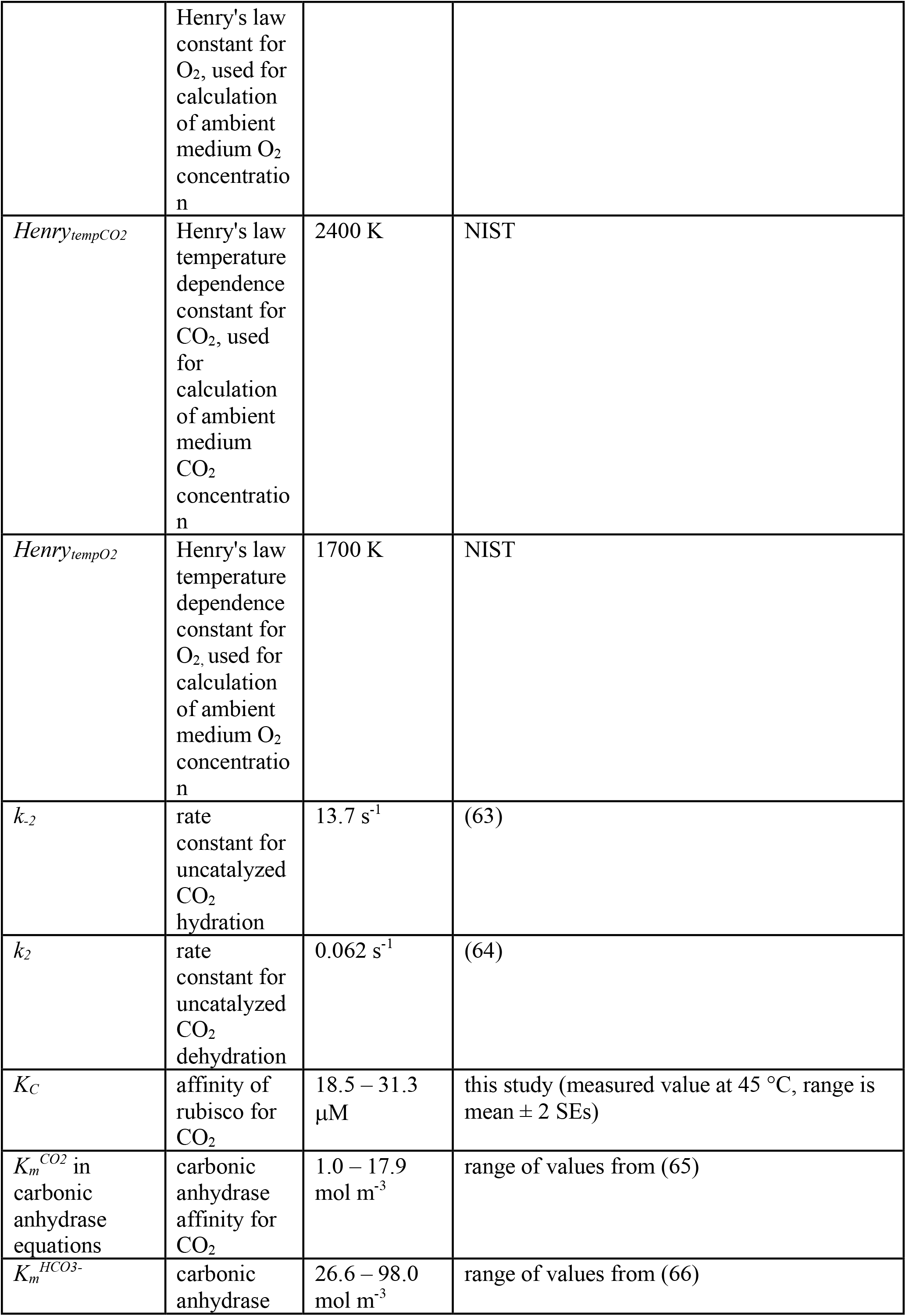
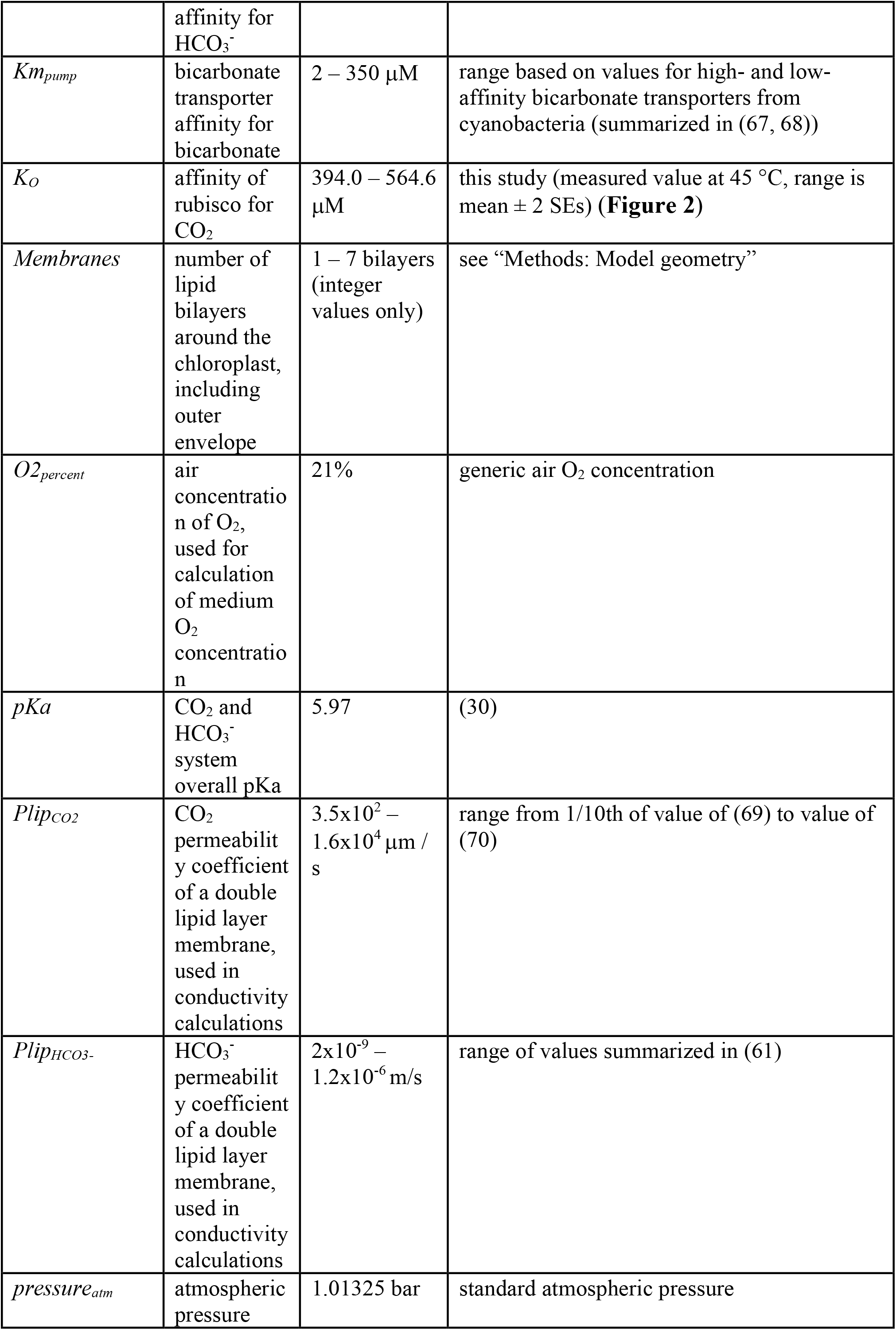
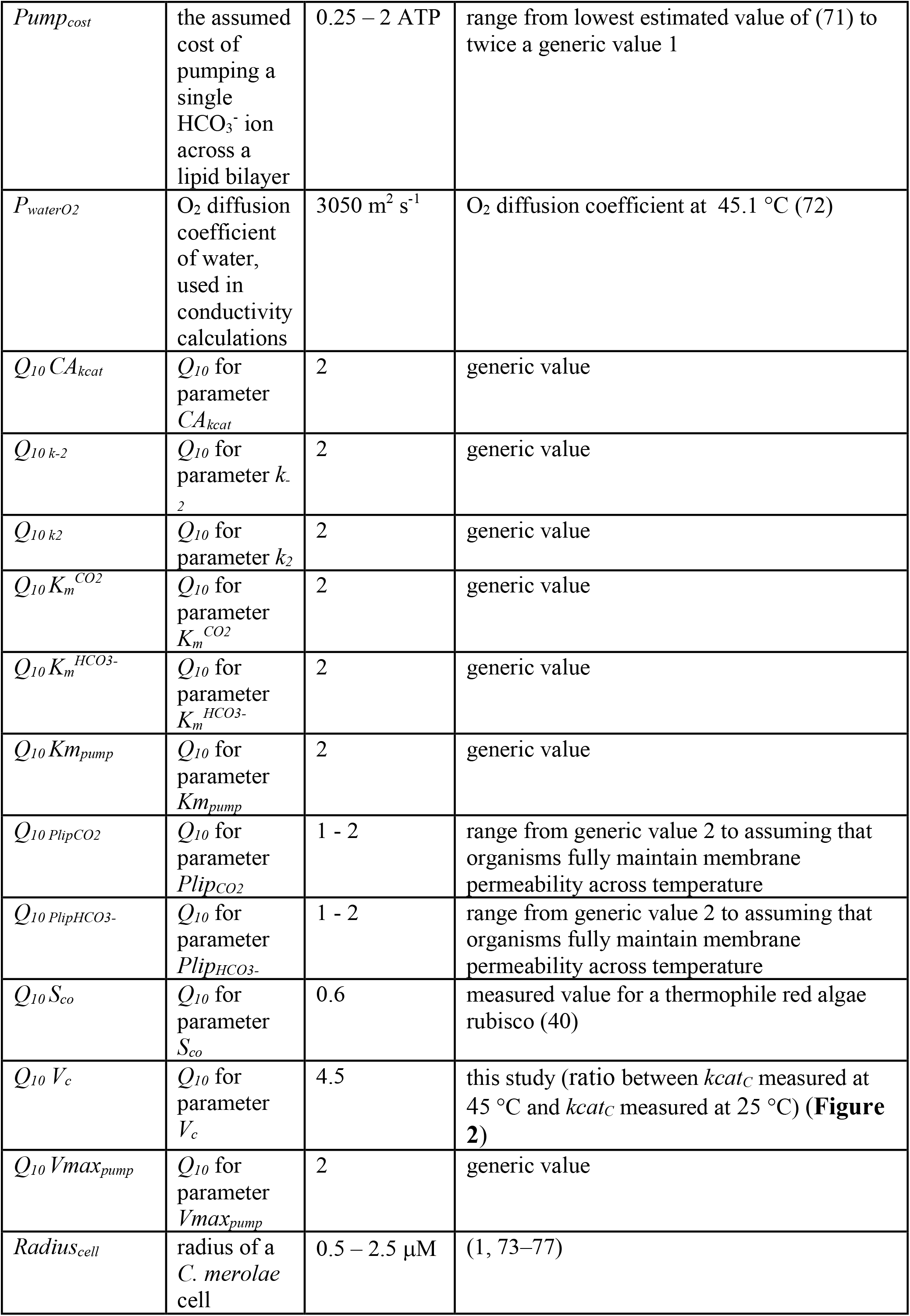
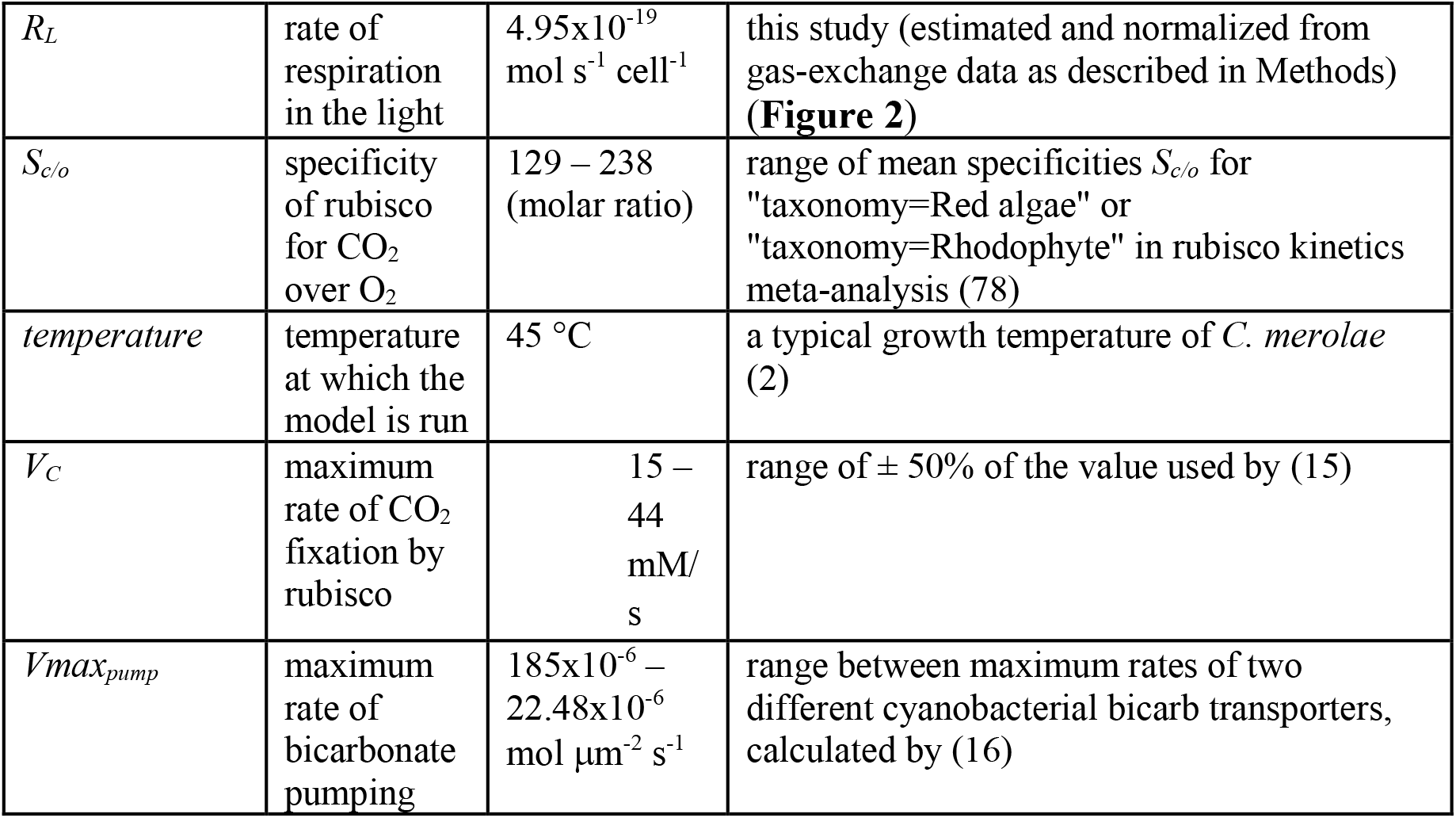
Parameter values or ranges used in the model. Values are known or assumed to be at 25 °C, unless otherwise specified.

**Table S2.** The test root-mean-square errors (RMSEs) and normalized RMSEs (NRMSEs) of four machine-learning surrogate models: eXtreme Gradient Boosting (XGBoost), Local approximate Gaussian Process (laGP), single-layer Neural Network (NN), and Deep Neural Network (DNN).

| **RMSE** | **XGBoost** | **laGP** | **NN** | **DNN** |
| --- | --- | --- | --- | --- |
| $\Gamma_{CO_2}$ | 0.1982 | 1.5054 | 0.7899 | 0.1458 |
| stromal CO<sub>2</sub> | 15.7500 | 73.2884 | 66.0004 | 15.6920 |
| $v_o/v_c$ | 0.0132 | 0.1248 | 0.0547 | 0.0613 |
| ATP per CO<sub>2</sub> | 49.8090 | 173.7715 | 161.0012 | 72.2325 |
| **NRMSE** | **XGBoost** | **laGP** | **NN** | **DNN** |
| $\Gamma_{CO_2}$ | 0.0142 | 0.1080 | 0.0567 | 0.0105 |
| stromal CO<sub>2</sub> | 0.0043 | 0.0201 | 0.0181 | 0.0043 |
| $v_o/v_c$ | 0.0086 | 0.0813 | 0.0356 | 0.0400 |
| ATP per CO<sub>2</sub> | 0.0084 | 0.0293 | 0.0271 | 0.0121 |

**Figure S1.**
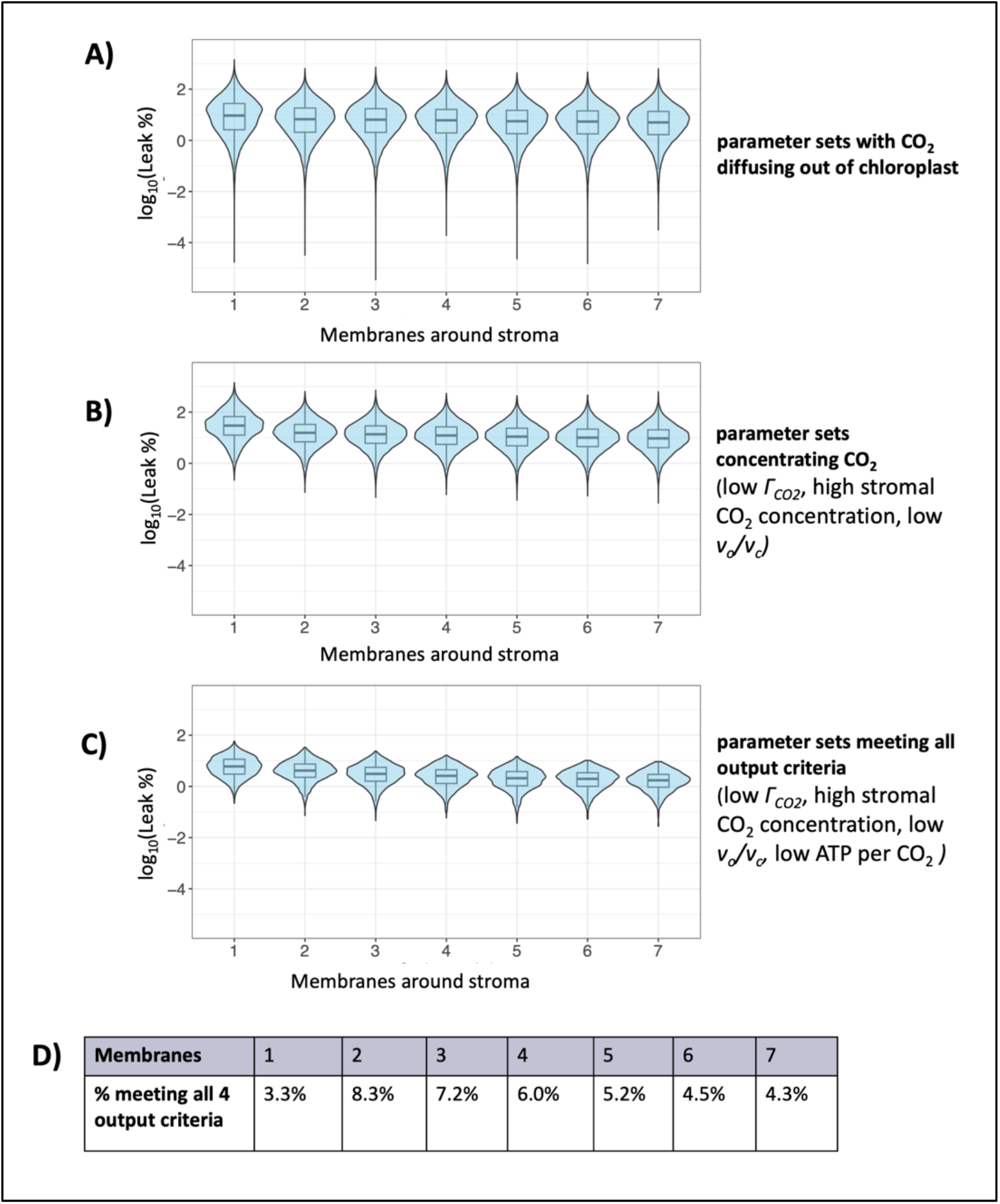
Effect of model input parameter *Membranes* (x-axis) on CO_2_ leakage from the chloroplast (represented as log_10_(Leak %): the log-transformed percentage relationship between the CO_2_ flux from the chloroplast to cytosol and the CO_2_ assimilation flux). **(A)** Results for parameter sets with CO_2_ diffusing out of the chloroplast (V5 steady-state flux towards cytosol, rather than towards chloroplast) (*n* = 191,345). **(B)** Results for parameter sets concentrating CO_2_ (low *Γ*_*CO2*_, high stromal CO_2_ concentration, low *v*_*o*_*/v*_*c*_) (*n* = 92,764). **(C)** Results for parameter sets meeting all output criteria (low *Γ*_*CO2*_, high stromal CO_2_ concentration, low *v*_*o*_*/v*_*c*_, low ATP per CO_2_) (*n* = 13,998).

**Figure S2.**
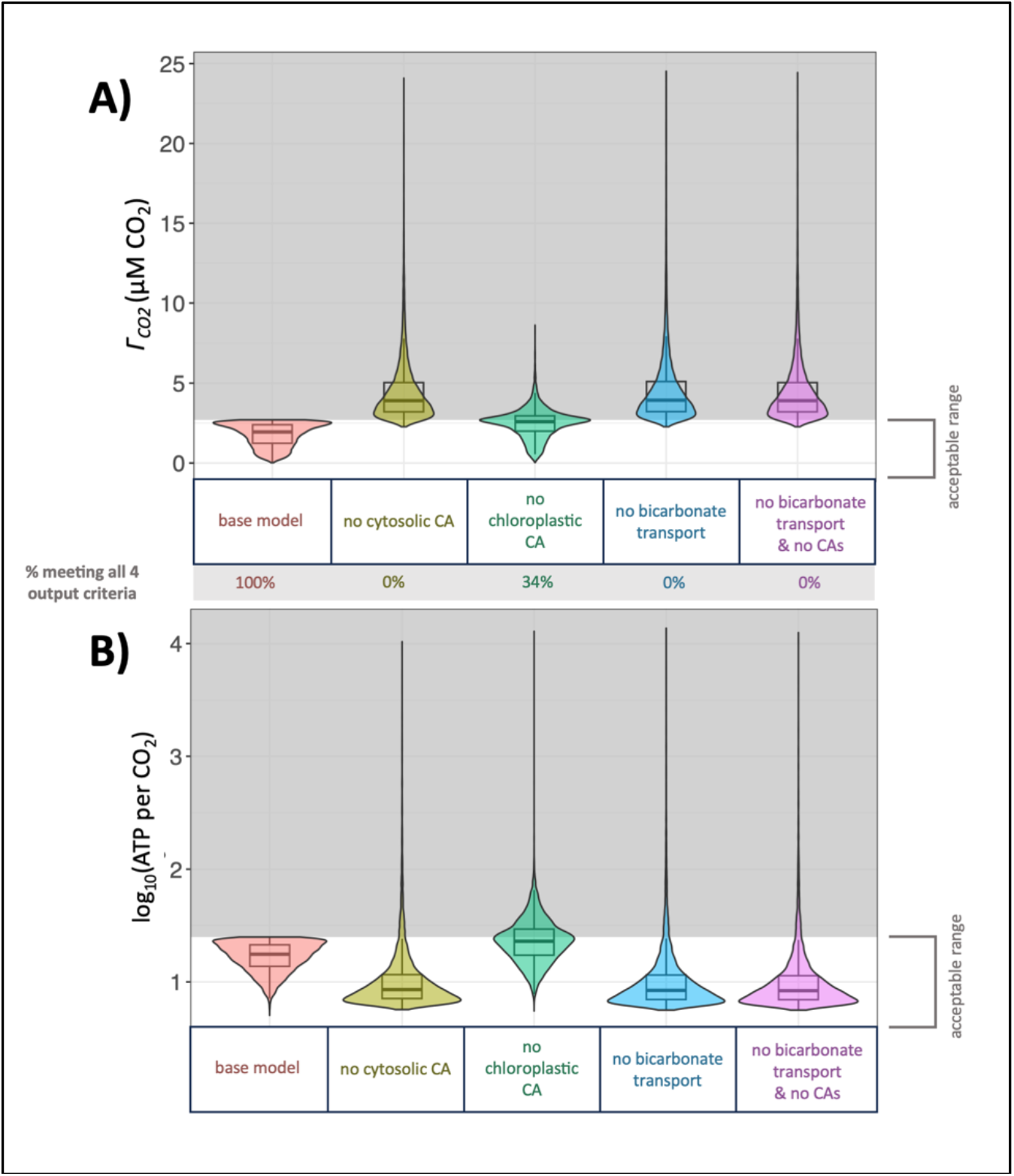
Effect on key model outputs when bicarbonate transport or carbonic anhydrases (CAs) are removed from the model, looking only at the 13,998 parameter sets that met all four output criteria in the base model. Distribution of parameter set outputs for each scenario is represented by a box plot overlaid on a violin plot. Shaded areas represent unacceptable values of outputs. The same sampling of input parameter sets was run through models representing each scenario. **(A)** *Γ*_*CO2*_ in model scenarios where various model features removed, with indication of how many parameter sets met output criteria in each scenario. **(B)** ATP per CO_2_ in model scenarios where bicarbonate transport activity at the chloroplast boundary is removed. 2,083 parameter sets producing negative ATP per CO_2_ values (3% of total) are not pictured in this panel due to log-transformation.

**Figure S3.**
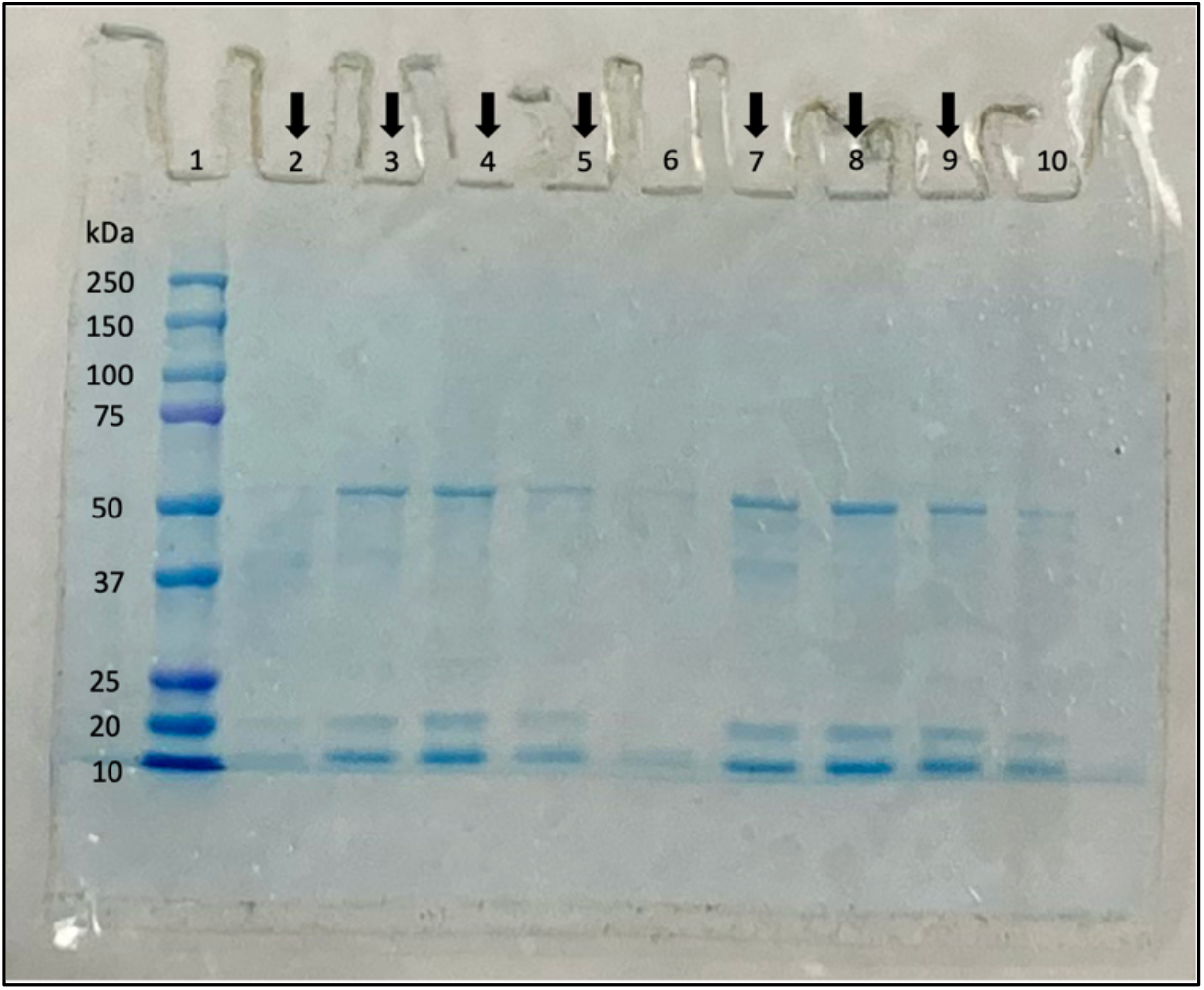
SDS-PAGE analysis of rubisco preparation. Lane 1: protein standards (BioRad Precision Plus Protein Dual Color Standards). Lanes 2 - 10 : fractions from fast protein liquid chromatography (FPLC) rubisco preparation. Three major bands are observable on the gel, which are believed to correspond to the rubisco large subunit (expected molecular weight 54 kDa), the rubisco small subunit (expected molecular weight 16 kDa), and the accessory pigment phycocyanin (composed of two subunits with reported molecular weights of about 15 – 18 kDa each). Arrows indicate fractions with rubisco activity detectable by a spectrophotometric assay, which were pooled for further preparation and analysis (see Methods for details). Molecular weights of rubisco were predicted from amino acid sequences CMV013C and CMV014C (*Cyanidioschyzon merolae* Genome Project v3, http://czon.jp/) (75, 79) using the Protein Molecular Weight Tool from the bioinformatics.org Sequence Manipulation Suite. Molecular weights of phycocyanin were reported by (4, 80).

**Figure S4.**
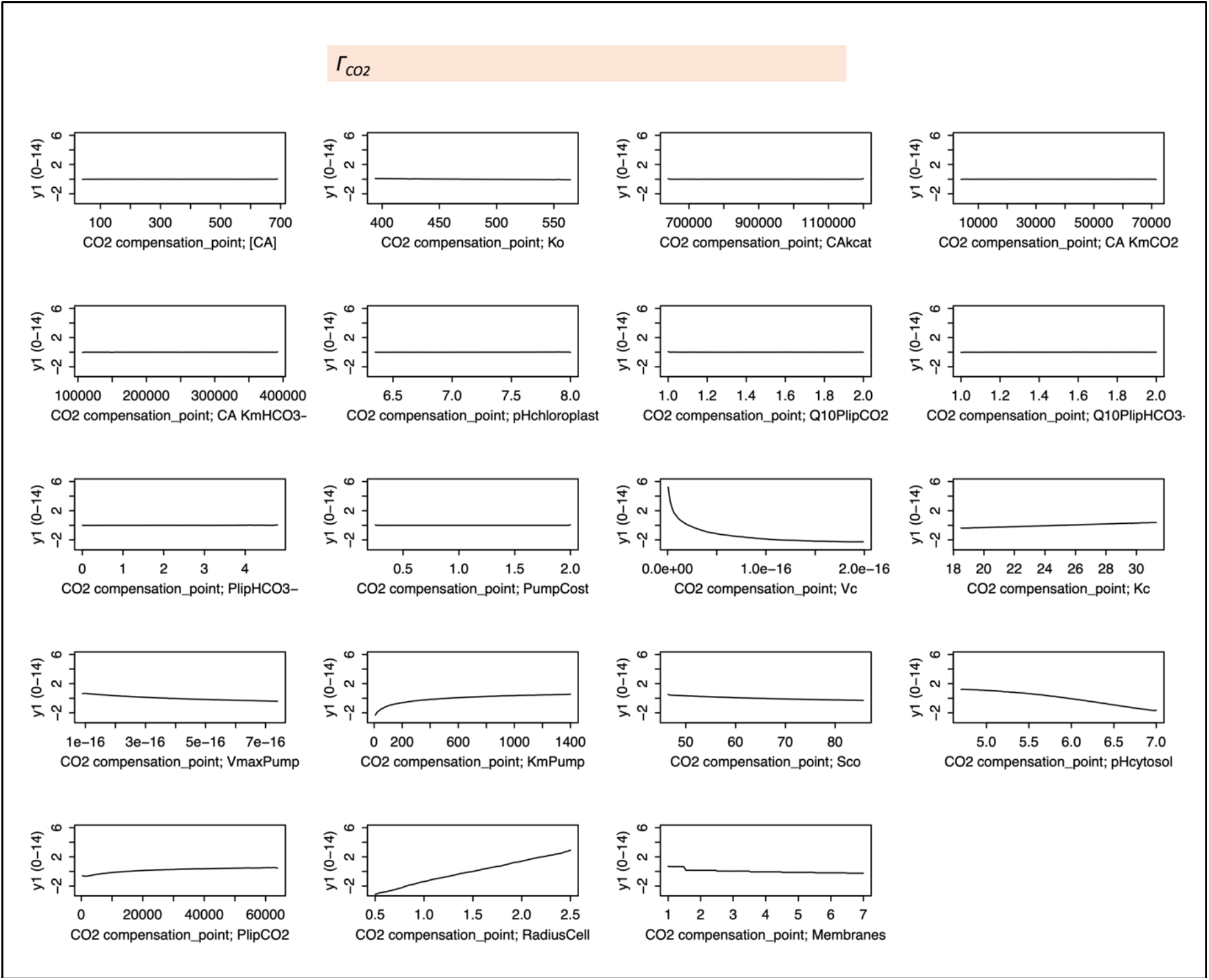
Partial dependence (PD) plots of first-order effects for *Γ*_*CO2*_.

**Figure S5.**
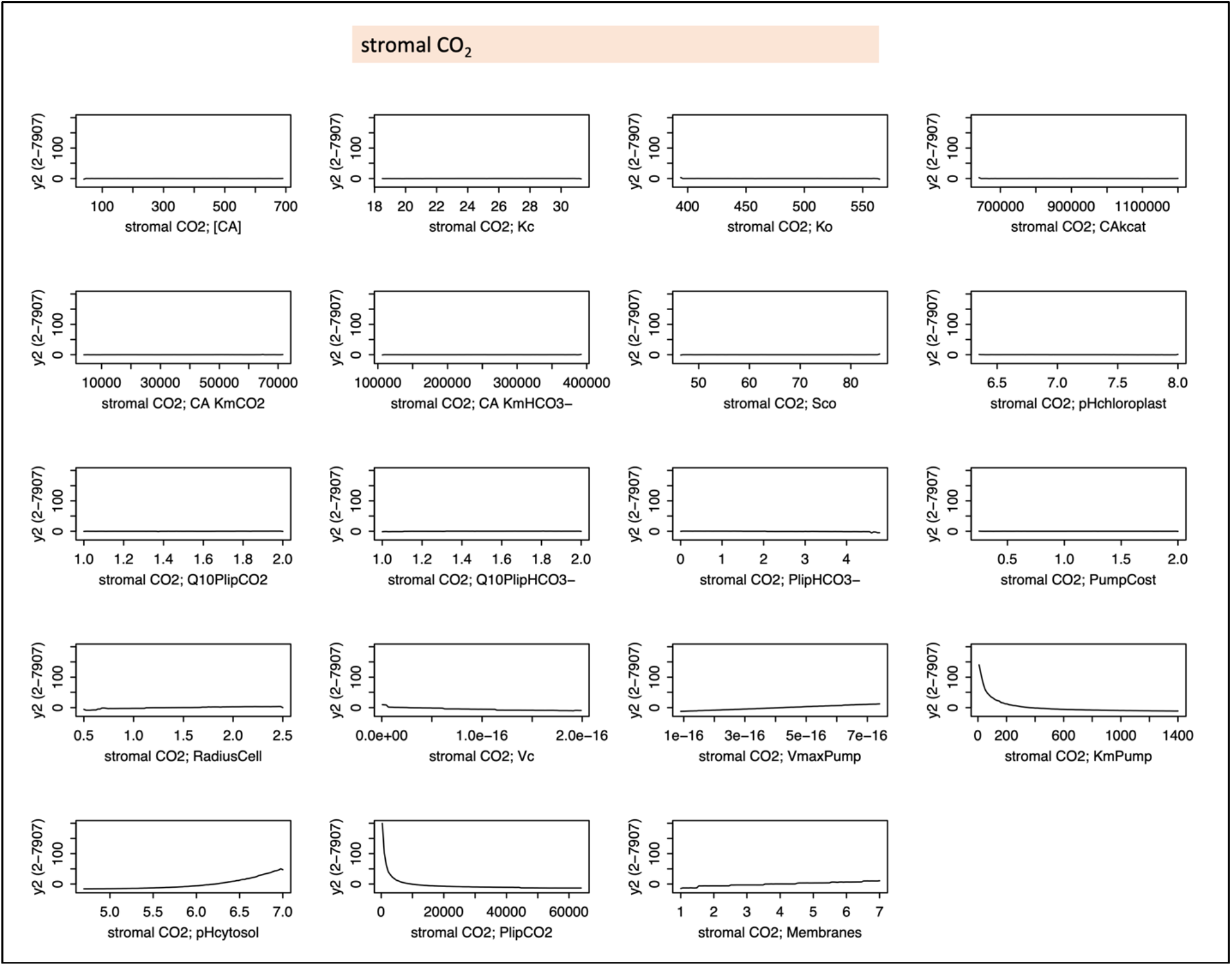
Partial dependence (PD) plots of first-order effects for stromal CO_2_.

**Figure S6.**
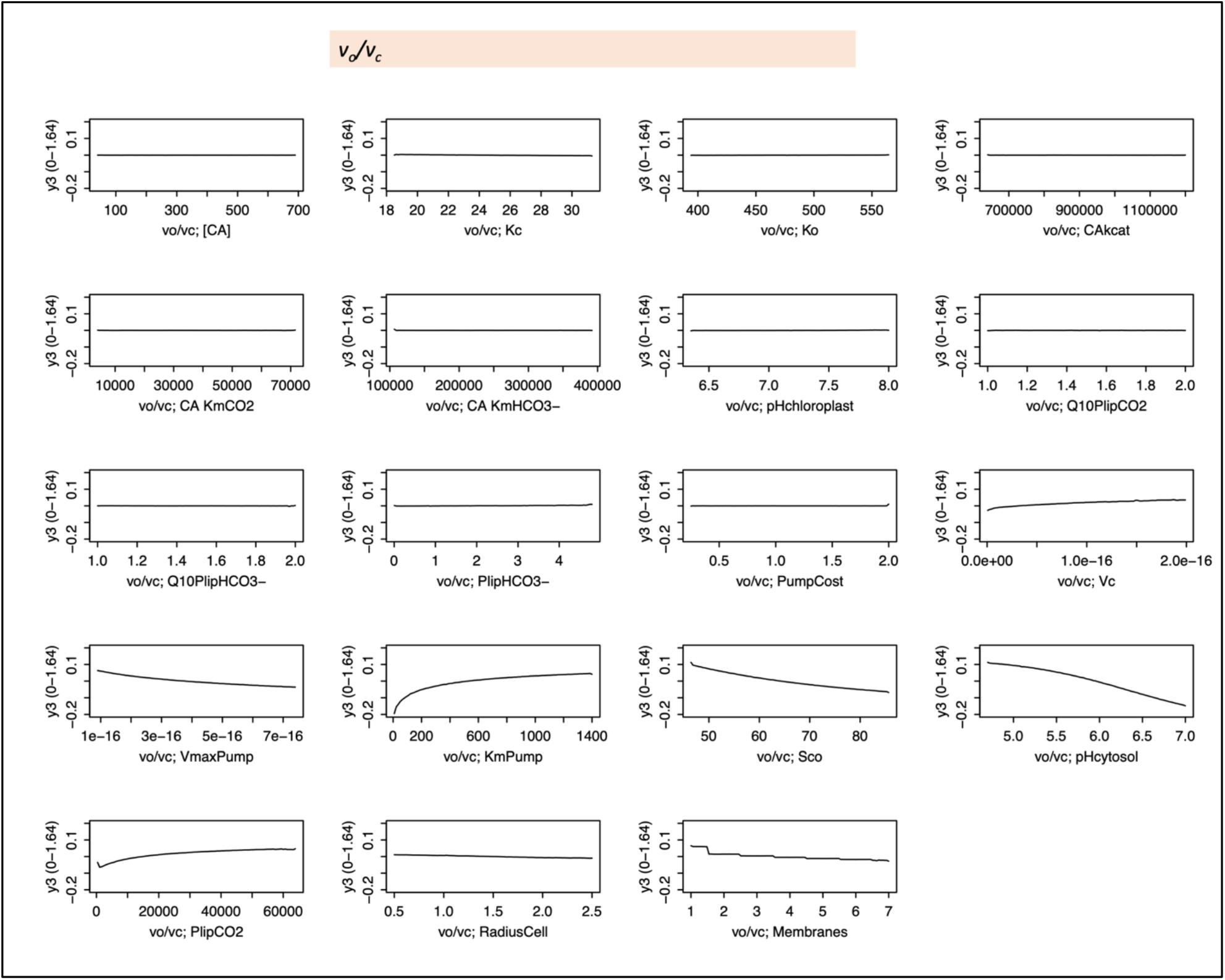
Partial dependence (PD) plots of first-order effects for *v*_*o*_*/v*_*c*_.

**Figure S7.**
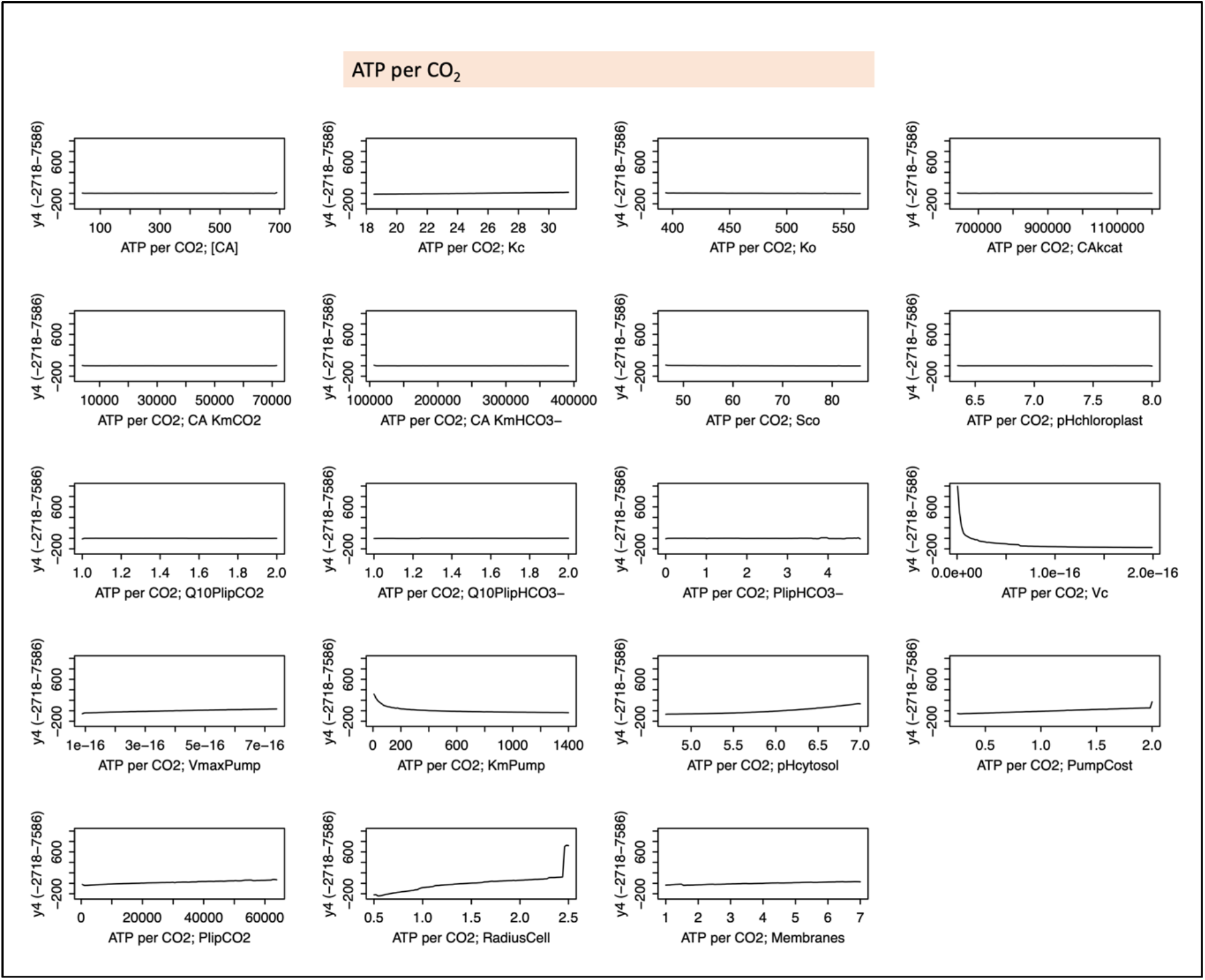
Partial dependence (PD) plots of first-order effects for ATP per CO_2_.

**Figure S8.**
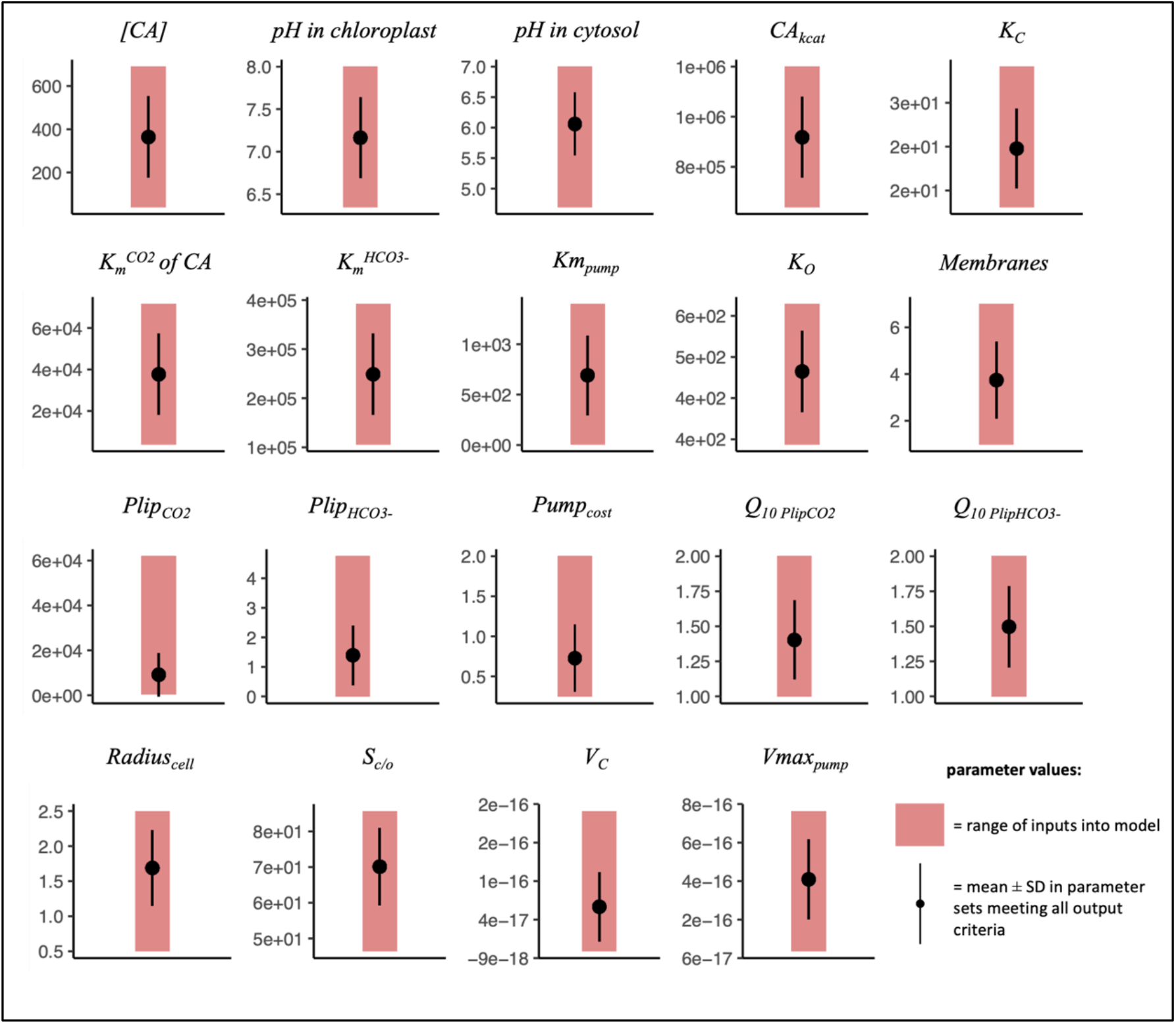
Ranges of parameters in all parameter sets (pink shaded areas) versus in parameter sets meeting all output criteria (black points with error bars indicating ± one standard deviation.

